# Non-coding mutations at enhancer clusters contribute to pancreatic ductal adenocarcinoma

**DOI:** 10.1101/2023.06.28.546873

**Authors:** Minal B. Patel, Eleni Maniati, Santosh S. Atanur, Debosree Pal, Ana Rio-Machin, James Heward, Hemant M. Kocher, Jude Fitzgibbon, Madapura M. Pradeepa, Jun Wang

## Abstract

Non-coding mutations (NCMs) that perturb the function of *cis*-regulatory elements (CRE, enhancers) contribute to cancer. Due to the vast search space, mutation abundance and indirect activity of non-coding sequences, it is challenging to identify which somatic NCMs are contributing to tumour development and progression. Here, we focus our investigation on the somatic NCMs that are enriched at enhancers from 659 pancreatic ductal adenocarcinoma (PDAC) tumours. We identify *cis*-regulatory NCMs within PDAC-specific enhancers derived from high and low-grade PDAC cell lines and patient derived organoids using two independent computational approaches. Five such CREs enriched for PDAC associated NCMs are also frequently mutated in other common solid tumours. Functional validation using STARR-seq reporter assays enables the prioritisation of 43 NCMs (7.3%) from a pool of 587 NCMs with 6,082 oligos, that significantly alter reporter enhancer activity compared to wild-type sequences. CRISPRi perturbation of an enhancer cluster harbouring NCMs over long non-coding RNA gene *MIR100HG*, which hosts a microRNA cluster (mir100-let7a-2-125b-1), leads to the downregulation of MIR100HG accompanied by a significant reduction in the TGF-β pathway (known to induce *MIR100HG*) and other PDAC critical pathways, including KRAS, p53, MTOR and TNF α signalling. Collectively, we have reported here *cis*-regulatory NCMs in PDAC proximal to many cancer-relevant genes, and our integrated approach paves way to explore CRE-associated NCMs in other human cancer genomes.

## Introduction

Pancreatic cancer, ranking fourth in the cause of cancer death in developed countries, is an aggressive malignancy with a devastating five-year survival rate below 9% after diagnosis ^1^. Pancreatic ductal adenocarcinoma (PDAC) is the predominant form of pancreatic cancer, encompassing approximately 90% of all cases^1^. Our understanding of the genomic landscape of PDAC is still mainly restricted to the somatic mutations within the coding regions of genes involved in PDAC ^2-6^. Our knowledge of non-coding mutations (NCMs) and their functional consequences in the development and progression of PDAC is still limited. The availability of large-scale whole genome sequencing (WGS) projects, such as those by the International Cancer Genome Consortium (ICGC) ^7^, along with assays profiling chromatin modifications, accessibility and conformation, has allowed for a systematic search for functional NCMs in various cancer types^8-14^.

Recent large-scale sequencing efforts by the Pan-Cancer Analysis of Whole Genomes (PCAWG) in over 2,600 primary tumours have identified several novel non-coding driver candidates, including NCMs in the 5’ region of *TP53* and 3’UTR o*N*f *FKBIZ* and *TOB1* using a statistically rigorous strategy for combining significance levels from multiple methods of driver discovery ^14^. More recently, Dietlein et al., implemented a genome-wide, sliding-window approach to detect significantly recurrent mutated regions across the whole genomes of 3,949 patients and 19 cancer types, considering chromatin features, tissue specificity and background mutations. Using this approach, they identified NCMs in CREs near canonical cancer genes and tissue-specific genes, such as regulatory regions proximal to *HIST1H1B* and *TMEM151A* in PDAC genomes and pancreas tissue-specific genes *CPB1* and *PNLIP* ^15^. Previously, Feigin et al., performed the PDAC-specific promoter-centric analysis and described Genomic Enrichment Computational Clustering Operation (GECCO) to uncover recurrent regulatory mutations in the *cis*-regulatory regions of 308 patient genomes. This method identified 16 genes with significant NCMs associated with promoter regions, and these genes were enriched for canonical PDAC pathways such as cell adhesion, axon guidance and Wnt signalling ^16^. However, previous methods have not fully or effectively utilised PDAC-specific epigenomic data in the discovery analysis, particularly active enhancer regions, leaving a large number of putative gene regulatory NCMs unexplored and PDAC-specific enhancer drivers unidentified. Such active enhancer-centric methods have previously been implemented in T-cell acute lymphoblastic leukaemia using sequencing reads derived from chromatin immunoprecipitation followed by sequencing (ChIP-seq) of histone H3 lysine 27 (H3K27ac) acetylation to identify recurrent enhancer associated variants^17^. Focusing on enhancer regions significantly reduces the non-coding genome search space to regions where non-coding variants are most likely to have potential functional activity at the gene control level^12,17,18^.

To address these specific challenges, we integrated epigenomic datasets for histone modifications associated with enhancers to identify PDAC-specific active enhancers and promoter regions. Together with gene expression profiles (GEP) where available and simple somatic mutation (SSM) data of 659 PDAC patients from ICGC, to investigate NCMs associated with PDAC-specific *cis*-regulatory elements (CRE). We implemented a composite of two independent approaches to detect putative CRE drivers enriched for significant NCMs. We further tested the regulatory activity of NCMs within these CREs using the high-throughput functional screening approach STARR-seq, followed by the analysis of one enhancer cluster using CRISPR-interference (CRISPRi) of CREs with NCMs (Fig. 1a). Our study combines a systematic computational analysis and experimental validation, identifying important CRE drivers involving PDAC-relevant genes. It also demonstrates a versatile workflow to investigate CRE-associated NCMs in other disease genomes.

**Fig. 1.**
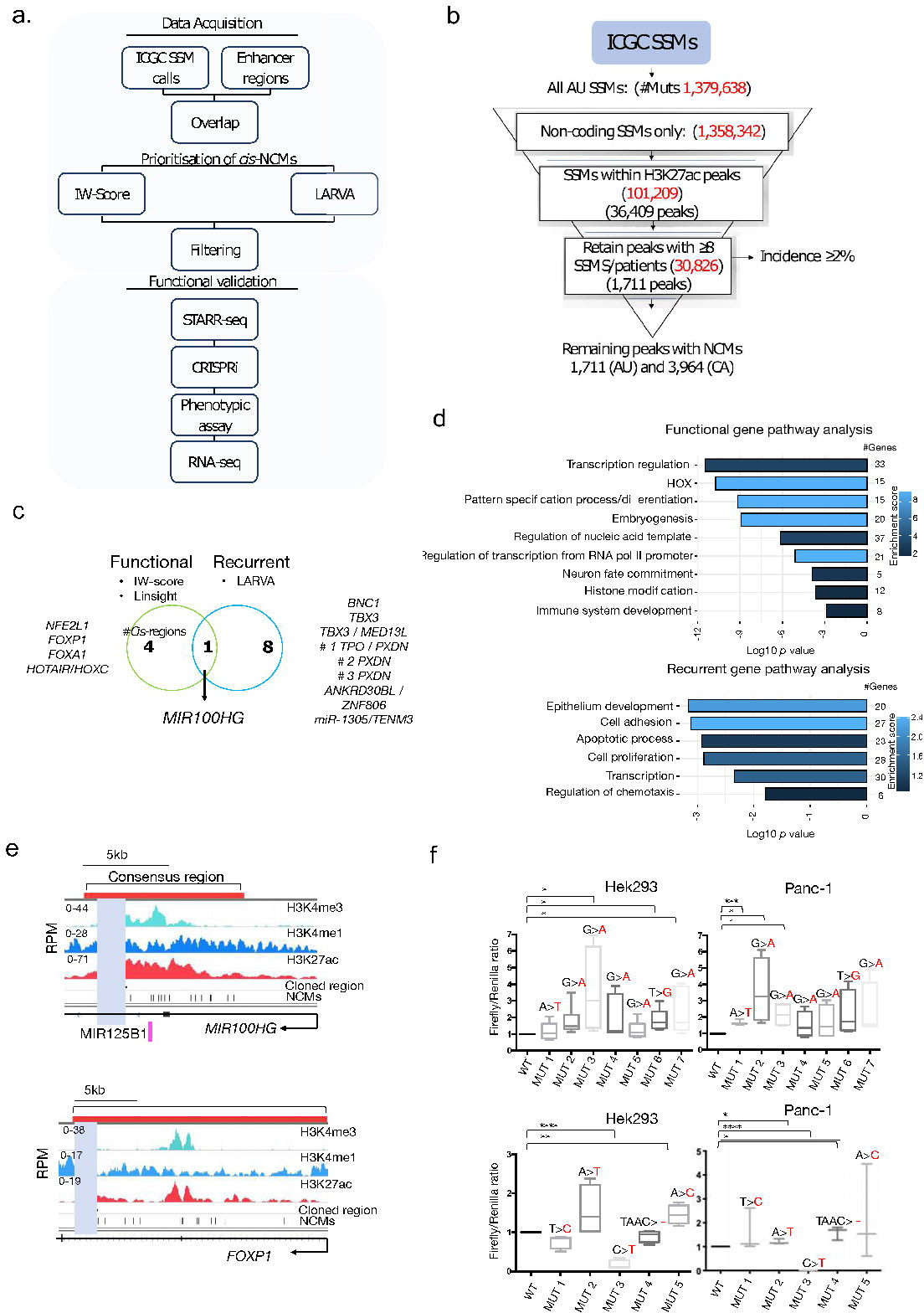
Identification of functionally significant PDAC-CRE-associated NCMs and putative CRE drivers in PDAC. **a.** Overview of our investigative strategy to detect significant CRE-associated NCMs and CRE drivers **b.** The variant filtering of somatic mutations using the ICGC PDAC Australia (AU) cohort as an example. The number of H3K27ac peaks and mutations (in red) is listed at each filtering step. **c.** Putative CRE drivers and the most proximal labelled genes identified by the two independent in-silico approaches: one implementing the IW-Scoring algorithm and LINSIGHT validation, the other using the LARVA model to identify CRE-regions with recurrent NCMs. **d.** Two gene set enrichment pathway analyses of CRE-associated nearby genes identified by the two in-silico approaches. **e.** Genome browser tracks (hg19) showing the histone modifications, CRE-associated NCMs in the third intron of *MIR100HG* and *FOXP1* selected for the Luciferase reporter assay validation (grey shade). **f.** Boxplots depicting the luciferase reporter activity of selected NCMs in the introns of *MIR100HG* and *FOXP1* tested in HEK293 and PANC-1 cell lines. The top panels are for NCMs in the *MIR100HG* CRE, and the bottom panels are for NCMs in the *FOXP1* CRE. Data is representative of 3 technical replicates from 3-4 independent experiments. The statistics was performed using the unpaired t-test, with the significance p-value shown as, *<0.05, **<0.01, ***<0.001, ****<0.0001.

## Results

### The mutational burden within PDAC *cis*-regulatory regions

To identify likely pathogenic NCMs in PDAC, we retrieved SSM data from the ICGC Pancreatic Cancer Genome Project Australia (AU, n=391 patients) and the Canada (CA, n=268 patients) cohort. 1,379,638 and 2,211,000 somatic mutations were identified in the AU and CA cohort, respectively. After filtering out non-synonymous somatic mutations, 1,358,342 (98.5%) and 2,179,517 (98.6%) somatic NCMs were retained from the AU and CA cohort, respectively, for further analysis. This corresponds to an average of 3,701 (AU) and 8,132 (CA) NCMs per patient.

We wanted to focus on the NCM burden within CREs, specifically those enriched with H3K27ac, a chromatin feature associated with active enhancers ^19^. We hypothesised that NCMs within these CREs may contribute to altering their function and target gene expression^17,20^. Using ChIP-seq datasets from seven PDAC cell lines ^21^ and two patient-derived organoid samples ^22^, we identified 404,415 enriched H3K27ac peaks across all samples. To consolidate H3K27ac peaks across the nine samples into one representative consensus region per loci, we stitched together quality peaks residing within 2,000bp of another (inter-peak distance), resulting in a total of 65,168 H3K27ac consensus peaks for further analysis (average peak length = 4,639 and SD = 7,949). This allowed us to narrow the search space for potentially important NCMs to ∼10% of the genome. Patient somatic NCMs were then mapped to the consensus H3K27ac coordinates to obtain a list of NCMs in PDAC-specific CREs. From the AU cohort of patients, 101,209 somatic mutations were observed within 36,409 (55.9%) consensus peaks and 166,541 CA cohort mutations within 43,002 (66.0%) consensus peaks. Therefore, capturing 7.45% and 7.64% of all AU and CA NCMs, respectively (Fig. 1b).

### Prioritisation of ***cis***-regulatory regions enriched with putative functional NCMs

We next aimed to interrogate somatic NCMs residing within consensus H3K27ac marked regions. To ensure the study of a significant proportion of PDAC patients, we retained CREs with a patient mutation incidence of 2% or above (n≥8), leaving 30,826 somatic mutations (AU cohort) across 1,711 consensus peaks/CREs and 64,867 somatic mutations (CA cohort) across 3,964 peaks (Fig. 1b). In total, 2.26% (AU) and 2.97% (CA) of the NCM burden remained to interrogate, similarly to observation in a previous study focusing on H3K27ac enriched elements^18^.

To prioritise the remaining CREs, we utilised two independent approaches: one measuring the functional effect of each NCM within a CRE and ranking them based on the median functional score of all NCMs; the other identifying CREs with significantly recurrent NCMs accounting for local background mutation rate, and replication timing (Fig. 1a). We carried out the first approach using the IW-scoring algorithm ^23^, an integrative weighted scoring framework to score NCMs and prioritised elements with a median IW-score of two or above (corresponding to a *p-*value ≤ 0.1). From the remaining 1,711 (AU) and 3,964 (CA) peaks after filtering, we identified 14 CREs from the AU-cohort and 32 elements in the CA-cohort using the median threshold (Extended Data Fig. 1 and S1). This method prioritised CREs annotated to cancer-related genes such as the *AP-1* transcription factor (TF) *JUNB* expressed in low-grade PDAC cells ^21,24^, and *GATA2,* associated with high-grade PDAC ^21^. Of the 46 prioritised CREs, five regions were shared between the AU and CA cohorts (Fig. 1c). These five CREs reside within the introns of the oncogenic long non-coding RNAs (lnRNA) *MIR100HG*^25^ and *HOTAIR*^26^ from and including the *HOXC* cluster of homeobox genes^27^, PDAC associated TFs *FOXA1*^28^ and *FOXP1*^29^ and ferroptosis related TF, *NFE2L1* ^30^ (Fig. 1c).

To further validate the putative significance of the NCMs within these five CREs, we compared the IW-score of NCMs residing within the H3K27ac positive regions to NCMs in immediate flanking sequences negative for H3K27ac marks. We observed a statistically significant higher IW-score of NCMs within H3K27ac enriched regions compared to those in flanking H3K27ac negative sequences (Extended Data S2), indicating the putative enhancer-associated NCMs have higher predicted functional consequences than mutations located outside these CREs. We also verified these findings with an independent scoring algorithm LINSIGHT, which scores variants on the likelihood of deleterious fitness consequences based on patterns of polymorphism and divergence from closely related species ^31^. The LINSIGHT model demonstrated a significant increase in the selective constraint (i.e., more deleterious on fitness) of H3K27ac-associated NCMs compared to NCMs in nearby H3K27ac negative regions (Extended Data Fig. 2).

**Fig. 2.**
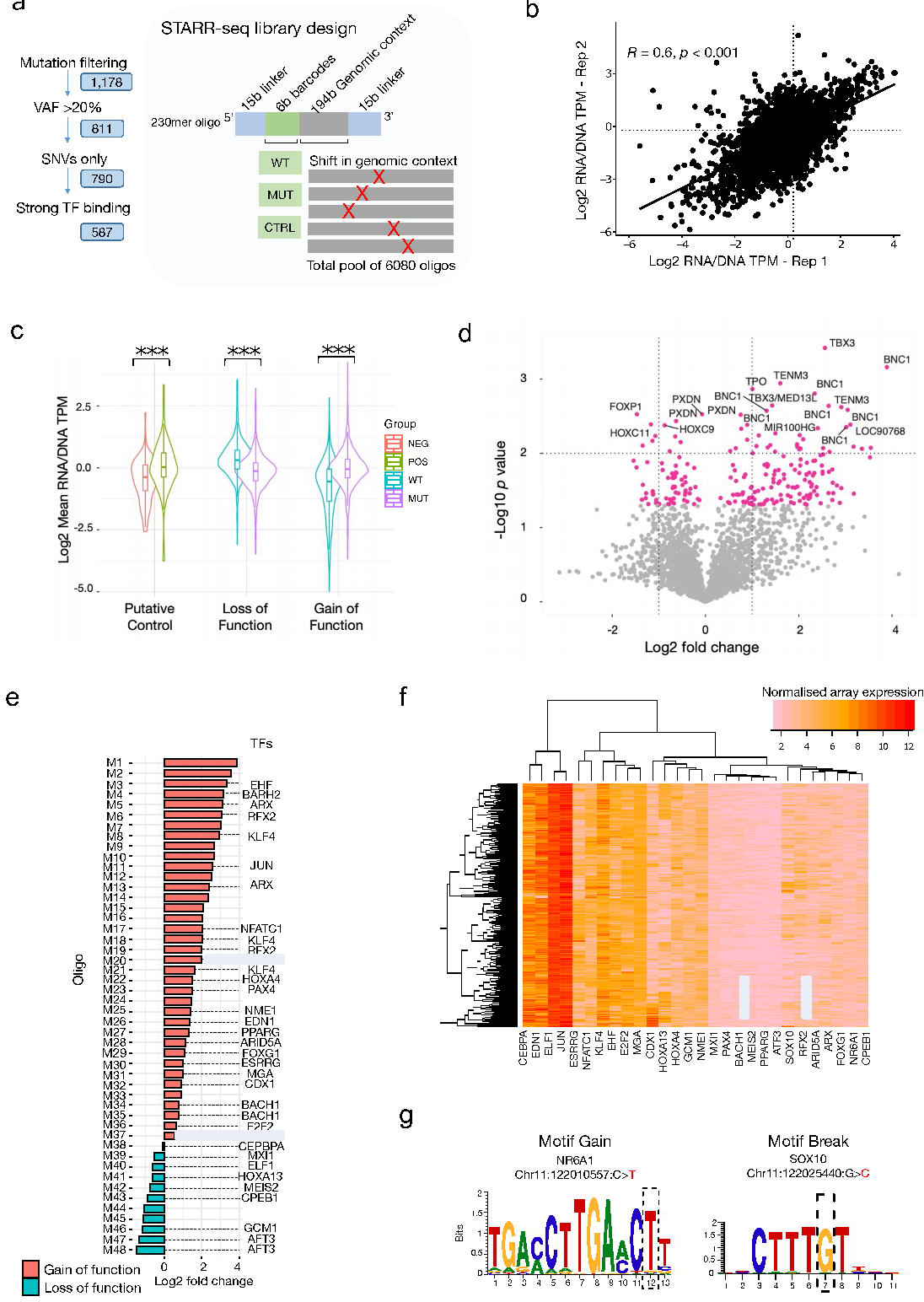
STARR-seq to validate the regulatory activity of candidate NCMs. **a.** STARR-seq NCM candidate selection strategy (left) and oligo design (right). **b.** A scatter plot showing the correlation of the STARR-seq regulatory activity between the two biological replicates. STARR-seq activity was measured as the log _2_ transformed transcript per million (TPM) of RNA output over the DNA input. The correlation coefficient (*R*) and *p*-values are shown. **c.** Violin plots depicting the mean log _2_ transformed STARR-seq activity (TPM) of the two replicates comparing the negative and positive controls, p***≤0.001 (t-test). **d.** Volcano plot showing the mean log_2_ fold change *vs.* the log10 p-value (t-test) between MUT and WT oligos for all constructs. Pink dots demonstrate candidates with a *p*-value <0.05. Selected candidate CREs with a *p*-value <0.01 (t-test) are labelled with the closest proximal gene. **e.** Oligos with the most significant changes compared to its WT counterpart (p<0.01). MUT oligos with a higher activity than their WT sequence (gain of function) are in red bars, while MUT oligos with a lower activity than the WT control are in green. Predicted motifs identified by MotifbreakR are shown beside bars for mutations where relevant. Oligo names M1-48 are listed in Extended Data S5. **f.** Heatmap showing the gene expression profile (GEP) from the ICGC PDAC cohort (n=269) of predicted TFs putatively perturbed or gained in the top significant NCMs (p <0.01). Normalised microarray expression values are shown in the heatmap. **g.** Motif gain and loss (break) from two mutations in the *MIR100HR* enhancer cluster. The TF binding motifs for TFs NR6A1 and SOX10 are shown, and the affected nucleotide is marked in a dotted line.

Using the second approach to identify significantly recurrently mutated CREs, we implemented LARVA ^32^. The LARVA model yielded 68 (AU cohort) and 71 (CA cohort) candidate CREs which were significantly recurrently mutated in relation to nearby background sequences (Benjamini-Hochberg (BH) adjusted *p* ≤0.01). These significant regions collectively harboured 1,842 and 2,258 NCMs in the AU and CA cohorts. Many NCMs were located proximally to several well-known genes implicated in PDAC, for example, an intergenic regulatory region in proximity to the miRNA: miR-21 and the Wnt/β-catenin signalling protein gene *WNT7b* (Extended Data S3). Nine significantly mutated CREs were shared between AU and CA cohorts. These recurrent CREs included regions proximal to the TF genes *TBX3* and *BNC1*, previously reported in PDAC^33,34^. NCMs were also located proximal to the adhesion molecule *PXDN*^35^ and transmembrane protein TENM3 ^36^, the lncRNA gene *TBX5-AS1*^37^, and microRNA, miR-1305 (Fig. 1c).

Notably, the *MIR100HG* enhancer cluster was the only one prioritised in the two approaches, but consisting of two separate CREs (Fig. 1c, Extended Data Fig. 3). Overall, our computational strategy has revealed NCMs enriched within or proximal to PDAC or cancer-related genes, including candidates identified from a previous non-coding study in PDAC^16^.

**Fig. 3.**
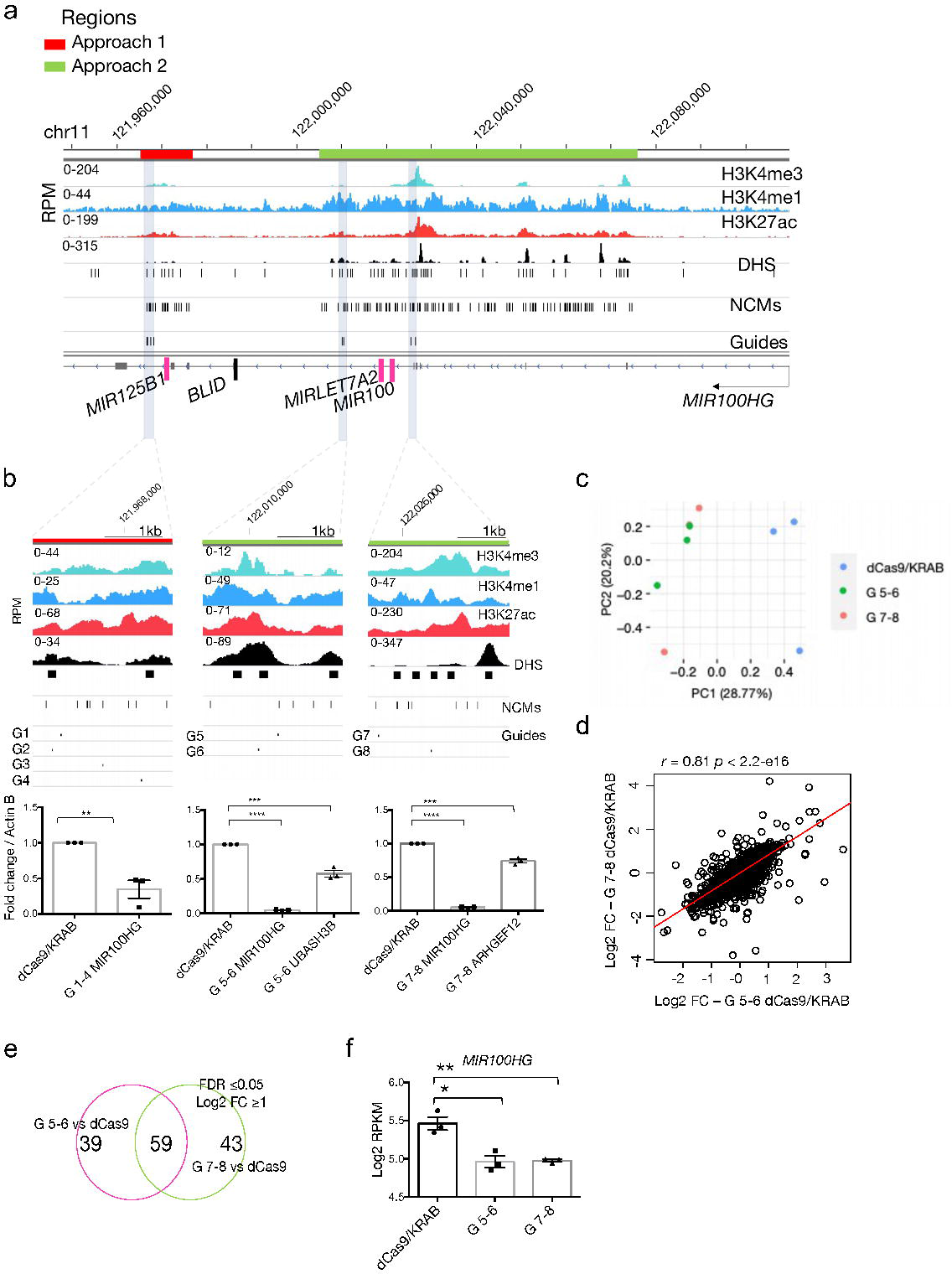
CRISPRi for selected CREs with NCMs within the MIR100HG enhancer cluster. **a.** Genome browser tracks (hg19) showing the overview of the *cis*-regulatory landscape at the MIR100HG enhancer cluster (11q24.1) and the selected CRE’s for CRISPRi perturbation (grey vertical bars). The first region (left) is within a significant CRE identified by the first *in-silico* approach based on IW-Scoring, and two regions (centre right) within the significant CRE by the second approach, based on LARVA. H3K27ac and H3K4me1/3, DNase I hypersensitive sites (DHS), NCMs, guide RNA sites, microRNAs and the *BLID* gene are shown. **b.** Zoom-in of the three targeted *MIR100HG* CRE regions. RTqPCR data showing fold change in MIR100HG, *UBASH3B* (for region two G5-6) and *ARHGEF12* (for region two G7-8) levels normalised to Actin-β upon CRISPRi compared to no guide RNA control (dCAS9/KRAB). **c.** P r i n c i p a l component analysis (PCA) of the RNA-seq samples among the three groups, dCas9/KRAB control, CRISPRi for region two (G 5-6) and region three (G 7-8). PC1 and PC2 were used for the separation of samples. **d.** Scatter plot of the lo_2_ gfold changes between G 5-6 and G 7-8 groups in comparison to the dCas9/KRAB control group for all profiled genes in the RNA-seq data. The correlation coefficient and *p*-value are shown. **e.** T h e o v e r l a p o f s i g n i f i c a n t l y differentially expressed (DE) genes between G 5-6 and G 7-8 groups in comparison to the control. The significance cut-off is shown, and numbers of shared and unique DE genes are listed. **f.** The level of gene expression of *MIR100HG* among the dCas9/KRAB, G 5-6 and G 7-8 groups were derived from the RNA-seq data (n=3 in each group). Log _2_ RPKM values were used to measure the RNA expression. A t-test was performed between the groups, with the significance p-value shown as *<0.05, **<0.01.

### Proximal genes to enhancer NCMs are associated with transcription and PDAC-linked biological processes

We next performed pathway enrichment analysis based on the annotated genes proximal to CREs identified by the two *in-silico* approaches using the DAVID tool ^38^. Inputting 95 annotated genes associated with 41 CREs identified by the IW-scoring approach, we observed significant enrichment in several gene families and regulatory processes, including homeobox genes, pattern specification, embryogenesis and transcriptional regulation pathways (Fig. 1d). Additional pathway analysis based on 212 genes annotated to the 130 recurrently mutated CREs identified significant enrichment in core molecular pathways including cell adhesion, epithelium development, cell proliferation, transcription, apoptotic processes and regulation of chemotaxis (Fig. 1d). The involvement of biological processes, such as embryogenesis, apoptosis and cell adhesion, has been reported in a previous genomic landscape study^39^. Furthermore, our findings complement Feign *et al.* in identifying NCMs significantly associated with homeobox genes and transcriptional regulation ^16^. Our results suggest a convergent mode for CRE-associated NCMs in relation to biologically relevant coding genes in PDAC.

### Enhancer NCMs show altered transcriptional reporter activity

To determine the effect of NCMs on the transcriptional regulatory activity, we performed luciferase-based enhancer reporter assays for a subset of NCMs. We selected twelve NCMs from two CREs identified from the first approach (IW score), comprising 11 single nucleotide variants (SNV) and a single 4bp deletion. Five SNVs were selected from the third intron of the *FOXP 1*gene, and seven NCMs in the third intron of the lncRNA *MIR100HG* (Fig. 1e). Interestingly, the 2kb region surrounding the seven NCMs at the *MIR100HG* l o c u s l a c k detectable H3K27ac and H3K4me1 marks in most of the cell lines, except those derived from high-grade PDAC cells PANC-1 and PT45P1, suggesting this putative active enhancer is specific to high-grade PDAC (Fig. 1e, Extended Data Fig. 3). Luciferase reporter assays were carried out in the high-grade PDAC cell line PANC-1 and easily transfectable cell line HEK293T. Within the *MIR100HG* CRE, NCMs (MUT 3, 6 and 7) and (MUT 1-3) showed significant increases in reporter activity in HEK293T and PANC-1 cells, respectively (Fig. 1f). Overall, all NCMs at this *MIR100HG* CRE showed an increase in luciferase activity compared to WT sequences, suggesting NCMs within this CRE are potentially gain-of-function, i.e., increase regulatory activity. The ∼2kb regulatory element surrounding five selected NCMs within the third intron of *FOXP1* was positive for H3K27ac marks in six PDAC cell lines (except for MIA-PaCa2 cells), and two patient-derived organoid samples (Extended Data Fig. 4). Among the five NCMs tested, two NCMs in HEK293T cells and three in PANC-1 cells significantly altered luciferase expression. Most notably, mutation 3 (chr3:71104908:C>T, IW-score = 5.20, *p* = 0.006, LINSIGHT score = 97.2%) significantly decreased reporter gene expression in both cell lines (Fig. 1f). Interestingly, all five NCMs within the *FOXP1* putative enhancer demonstrated concordance in the overall transcriptional regulatory activity in both cell lines.

### STARR-seq assays prioritise a subset of 43 NCM candidates for further validation

Next, we screened a larger set of NCMs within consensus CREs using the high-throughput approach, Self-Transcribing Active Regulatory Region sequencing (STARR-seq)^40^. To focus on NCMs with the strongest evidence of predicted function, we retained 504 NCMs with a variant allele frequency above 20% and strong TF binding strength as predicted by motifbreakR^41^. Of the 504 NCMs, binding motifs of 258 TFs were strongly predicted to occupy these mutation sites. Moreover, among the 73 NCMs identified by the first approach (IW score), 47 (64%) NCMs were predicted to cause TF-motif gain and 26 (36%) loss-of-motif (break). Among the 431 NCMs selected from the second approach (LARVA), 216 (50%) NCMs caused predictive gain and 215 (50%) loss of motif changes. We included 83 single base indels, resulting in 587 candidate NCMs in the final STARR-seq library (Fig. 2a).

We designed ten 230bp oligos per NCM, five for each NCM and five for the corresponding wild type (WT). One oligo represented the NCM in the middle and four oligos had a 10 bp sliding genomic window (SW) in either direction from the centre of the oligo (Fig. 2a, see Methods). A further 210 positive (PDAC enhancers) and negative (no enhancer features) control oligos were included in the library, resulting in a pool of 6,082 oligos.

Sequencing and quality analysis of the cloned STARR-seq plasmid library demonstrated good complexity and accuracy (Extended Data Fig. 5), with comparable outcomes to a previous MPRA study using synthetically designed oligos ^42,43^. We performed two biological replicates of STARR-seq by transfecting the PANC-1 cell line (see methods)^44^. After filtering low-quality reads across samples, we observed a good concordance between replicates (Fig. 2b). As expected, positive control sequences showed significantly higher reporter activity compared to negative controls (Fig. 2c).

We next tested the significance between mutant (MUT) and WT constructs on reporter gene expression across replicates. A total of 217 plasmids (representing 155 NCMs) showed significant differential enhancer activity (log_2_ fold change −1.54 to 3.53, Student’s t-test, *p*<0.05). 95 (61.3%) NMCs showed significantly increased enhancer activity, while 60 (38.7%) mutations showed a significant reduction in enhancer activity in comparison to WT sequences (Fig. 2c and 2d). Interestingly, 36 CREs harbouring indels showed significant fold changes at similar activity to SNVs (mean log_2_ FC 1.07). Despite the differences in assays and genomic context, we observed concurrent directional changes in enhancer activity at NCMs assayed by luciferase reporter assays and sequencing-based high-throughput STARR-seq (Extended Data Fig. 6).

Focusing on the most significant alterations between MUT and WT alleles (t-test, *p*<0.01), we highlighted 43 mutations, 33 of which demonstrated an increase in reporter activity and 10 with an observed reduction (Fig. 2e). Notably, the differential activity changes between MUT and WT in 13 NCMs were significantly altered in three or more independent STARRs-seq plasmids (*p*<0.05). Similarly, 31 NCMs were significantly altered in two independent SWs demonstrating concurrent directional activity changes. Eight of the 43 NCMs were located within an enhancer cluster (observed in low-grade and MiaPaCa2 cells) upstream of the *BNC1* gene (Extended Data Fig. 7a). The NCMs proximal to *BNC1* significantly increased reporter gene expression in PANC-1 cells in comparison to WT sequences (Fig. 2d). Assessing the expression of genes within 1Mb of this consensus peak by comparing MUT and WT patient GEPs, we did not observe a difference in the expression of BNC1, previously reported to be methylated in early stage PDAC patients ^45^. However, we observed a significant increase in the expression of nearby genes *BTBD1* (*p* = 0.003, ∼234kb from the middle of the consensus peak to BTBD1 TSS), important in cell survival, the ubiquitin/proteosome degradation pathway and mesenchymal differentiation ^46^ and *FAM103A1* (*p* = 0.008, ∼316kb) which encodes an important subunit for the 7-methylguanosine cap added to the 5’ end of mRNA and an essential component for gene expression^47^. Patient GEP analysis also revealed a significant decrease in the transmembrane protein *TM6SF1* (frequently hypermethylated ^48,49^, *p* = 0.0003, ∼137kb) between MUT and WT patients (AU cohort, Extended Data Fig. 7b), overall suggesting these NCMs may exert their regulatory potential in a more distal manner. Additional significant increases in reporter gene expression were observed proximal to the PDAC-associated TF *TBX*^50^ (7 NCMs) and in the introns of lncRNA *MIR100HG* ^25^ (3 NCMS, Fig. 2d).

To assess the putative biological implications of these top-performing STARR-seq NCMs, we took a closer look at the TF-motif binding predictions. From the 35 NCMs in the top 43 STARR-seq performing mutations with TF-motif predictions, 21 were characterised as TF binding motif-gain (creating *de novo* TF binding motifs), while 14 were TF binding motif-loss. For example, one gain-of-function NCM proximal to *TBX3* (chr12:115067012:C>A) was predicted to create a binding motif for the oncogenic TF JUN (Fig. 2d and Extended Data Fig. 7c). This NCM led to a mean log_2_ fold-change of 3.69 in STARR-seq reporter gene expression across all five SWs (Mann Whitney U test, *p*=0.016). As expected, JUN was highly expressed in PDAC patients based on the patient GEP in the AU cohort (Fig. 2f) ^51^. The most significant loss-of-function was observed in a NCM located in the intron of *FOXP1* (chr3:71123616:G>T) supported by three significant SWs (*p* <0.05, average lo2gfold change across SWs = −1.36). At this site, the binding motif of an unfolded protein response (UPR) mediating TF, the activating TF-3 (ATF3) ^52^, was predicted to be disrupted (Extended Data Fig. 7d) and was found to be moderately expressed in the PDAC patient GEP (AU cohort, Fig. 2f). Furthermore, the top two NCMs located in the *MIR100HG* enhancer cluster also showed strong effects on TF binding: the first mutation (chr11:122010557:C>T) demonstrated a gain of TF motif, creating a *de novo* binding motif for *NR6A1*, a nuclear receptor family member; while the second mutation (chr11:122025440:G>C) was predicted to disrupt the binding motif for *SOX10* (Fig. 2g), a reported tumour suppressor through the suppression of the Wnt/β-catenin pathway in digestive cancers ^53^. We observed that *NR6A1* and *SOX10* TFs were expressed at moderate levels in PDAC patients (AU cohort, Fig. 2f). Overall, using the STARR-seq assay enabled the prioritisation of CRE-associated NCMs for further investigation.

### CRE cluster harbouring NCMs located at the *MIR100HG* locus regulates genes *in cis*

The two computational approaches used in this study identified the lncRNA MIR100HG locus as a significant candidate for harbouring NCMs in separate CREs in each approach. Notably, *MIR100HG* is host to the oncogenic miR-s pre-miR125b-1 and pre-miR-100, previously implicated in PDAC^25,54^, and they modulate (including *MIR100HG*) in a pro or anti-tumourigenic manner depending on the cancer ^25,54-59^. It hosts the tumour suppressors pre-miR-Let7a-2^25^ and the pro-apoptotic protein *BLID*^60^, located within intron three of MIR100HG (Fig. 3a).

Next, we investigated the functionality of three CREs harbouring NCMs at the *MIR100HG* enhancer cluster using a CRISPRi approach recruiting the dCAS9/KRAB repressor to NCMs and CREs of interest ^61^ (Fig. 3a and b). The first region located ∼2kb away from the hosted pre-miR-125b-1 in the third intron of *MIR100HG* harboured NCMs identified from the first *in silico* approach. CRISPRi with a pool of four independent lentiviral guide RNAs (G 1-4) were selected close to NCMs that were shown to alter enhancer activity in either luciferase or STARR-seq experiments (Fig. 1e, 3b). Two guides (G 5-6) were designed to target region two harbouring five NCMs (CRE-two), including the most significant NCM identified to drive reporter enhancer activity using STARR-seq (M20 in Fig. 2e and Fig. 3b). An additional two guides (G 7-8) were designed to target the third region harbouring six NCMs (CRE-three), including a gain-of-function NCM from the most significant STARR-seq candidates (M37 in Fig. 2e).

CRISPRi, followed by RT-qPCR, showed a significant reduction in MIR100HG expression in all three CREs in this enhancer cluster in comparison to dCAS9/KRAB negative controls (Fig. 3b). This data suggests that these CREs function as active enhancers to r e g u l a t e t h e e x p r*M*e*IR*s*10*s*0*i*H*o*G*.nAnoalyfsis of looping interactions from the 4D genome^62^ and integrated method for predicting enhancer targets (IM-PET) ^63^ data in PANC cells indicated interactions between CRE-two and the promoter of *UBASH3B* l o c a t e d upstream of *MIR100HG* (Extended Data Fig. 8a) *. UBASH3B* has been reported to inhibit the endocytosis of the epidermal growth factor (EGFR), an essential component in the development of pancreatic precursor lesions ^64-66^. RT-qPCR analysis demonstrated a significant decrease in *UBASH3B* expression with the CRE-two CRISPRi compared to controls (Fig. 3b). CRE-three shows interactions with the promoter of *ARHGEF12* (Extended Data Fig. 8b). *ARHGEF12,* a guanine nucleotide exchange factor (GEF), activates Rho A, a key regulator of cytoskeleton organisation and ROCK1/2 induced extracellular matrix remodelling, associated with poor outcomes in PDAC patients ^67^. CRE-three CRISPRi resulted in a significant decrease in *ARHGEF12* levels compared to controls (Fig. 3b). These results suggest that CRISPRi-based perturbation of CRE-two and three leads to downregulation of genes located *in cis*, although to a less extent compared to the reduction in *MIR100HG* expression.

### CRISPRi perturbation of MIR100HG CREs alters core PDAC signalling pathways and cell motility

We performed RNA-seq to evaluate the global mRNA changes in CRISPRi-targeted CRE-two and -three clones (Fig. 3b). Principle component and correlation analyses showed CRISPRi of CRE-two and -three shared similar gene expression programmes (Fig. 3c and 3d). Differential expression (DE) analysis identified 98 and 102 significant genes in the perturbation of CRE-two and -three clones compared to the control, respectively (FDR<0.05 and absolute log _2_ FC >1). Of them, 59 DE genes were shared between the two clones (Fig. 3e). We also observed a significant reduction in *MIR100HG* RNA-seq expression in both targeted CREs, consistent with the qPCR data (Fig. 3f).

Gene set enrichment analysis (GSEA)^68^ against the MsigDB Hallmark ^69^ and oncogenic signature gene sets were then performed between the two CRISPRi groups and the dCas9-KRAB control (Fig. 4a). In both CRISPRi perturbations; we observed a comparable and significant downregulation of important PDAC hallmark gene sets involved in KRAS signalling^70^, UPR, reactive oxygen species (ROS) ^71^ and T NαFsignalling ^72^ (Fig. 4a and 4b). Oncogenic signatures associated with critical drivers KRAS^73^, P53, epithelial-to-mesenchymal transition (EMT) inducing TGF-β and cell survival and proliferation-related MTOR ^73^ pathway genes were significantly reduced in both inhibited *cis*-regions. In contrast, migration inhibiting cAMP ^74^ and interestingly pro-EMT related LEF1 ^75^ signatures were significantly upregulated (Fig. 4a). Collectively, the CRISPRi perturbation of two CREs at MIR100HG led to a significant reduction in key oncogenic molecular mechanisms observed in PDAC, resulting in a more favourable phenotype.

**Fig. 4.**
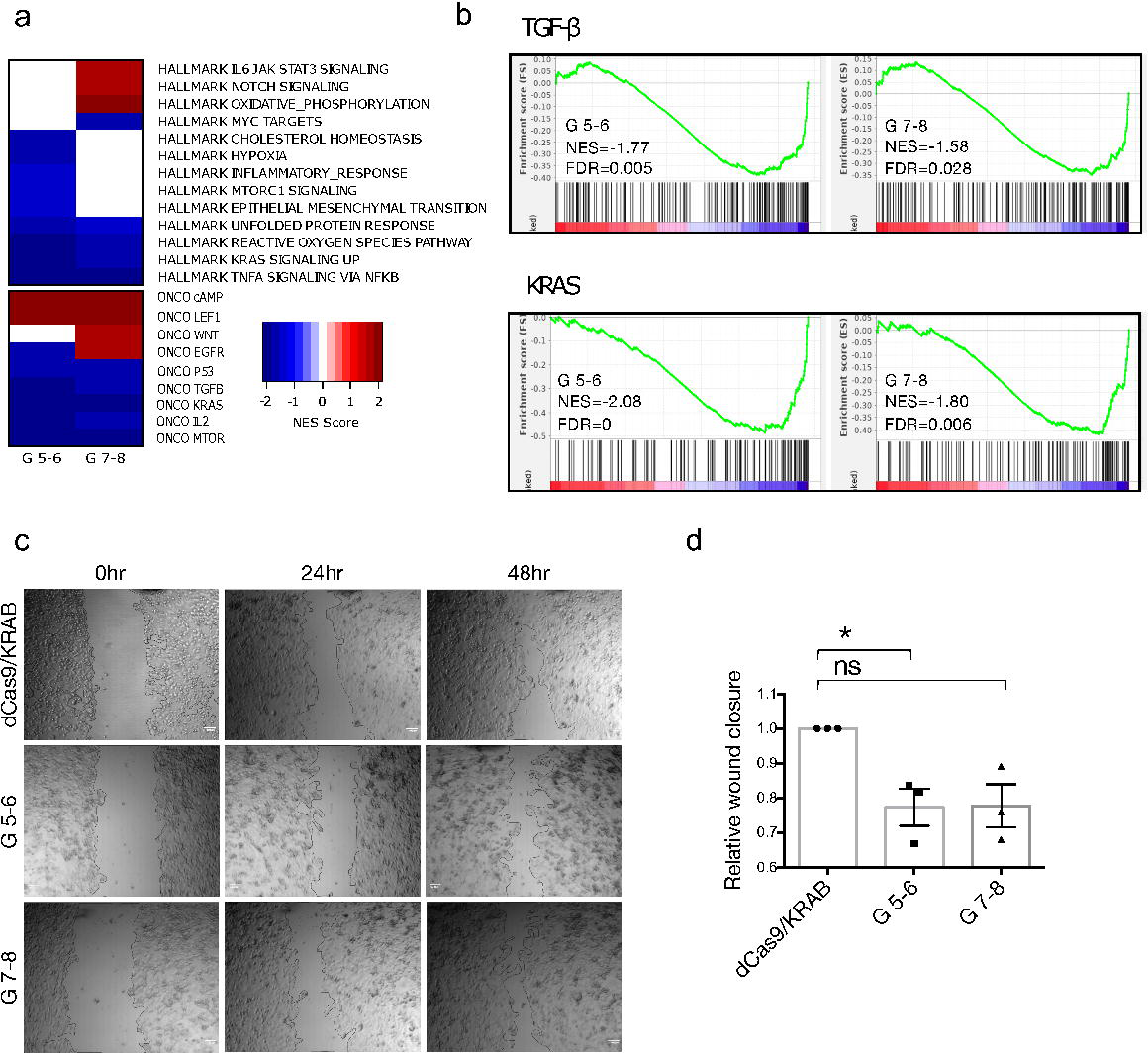
CRISPRi for *MIR100HG* CREs results in a downregulation of KRAS and TGF-β pathways. **a.** Significantly dysregulated pathways (false discovery rate, FDR<0.05) in the CRISPRi perturbation groups G 5-6 and G 7-8 compared to the dCas9/KRAB control group. Gene set enrichment analysis (GSEA) against the hallmark and oncogenic signature gene sets was performed ^68,69^. The normalised enrichment scores (NES) were used to create the heatmap, with the positive NESs (in red) indicating the upregulation and negative NESs (in blue) indicating the downregulation of activities in the CRISPRi perturbation groups compared to the dCas9/KRAB control. **b.** GSEA plots for the TGF-β and KRAS signalling gene sets for the CRISPRi perturbation G 5-6 and G 7-8 groups compared to the dCas9/KRAB control group. The NES and FDR values for each analysis are shown. **c.** Wound healing assay with G 5-6 and G 7-8 CRISPRi clones compared to the dCas9/KRAB control samples. 0, 24 and 48-hour time points are shown. **d.** Measurement of the relative wound closure in the three groups, dCas9/KRAB control, region two (G 5-6) and region three (G 7-8) (images are a representation of n=3 biological replicates in each group). An unpaired t-test was used to compare perturbation clones vs. control groups. p-values *≤0.05. ns, not significant.

TGF-β regulates *MIR100HG* transcription and thus the release of its hosted miRs, inducing EMT, encouraging cell motility and metastasis ^25^. Here, we identified many TGF -β related genes such as *FGF1*^76^, *KDM6B*^77^, *LIF*^78^, *PIK3CD*^79^, *PXDC1* a n d*TAGLN*^80^ w e r e significantly downregulated in the two CRISPRi groups compared to the control (Extended Data Fig. 9). Hence, we further aimed to validate the reduction in TGF-β signalling observed with GSEA enrichment by using wound healing assays (Fig. 4c). Over 48-hours, the inhibition of CRE-two (G 5-6) resulted in a significant reduction in cell motility in comparison to controls, corroborating with a stronger gene enrichment reduction in TGF-β and EMT signalling compared to CRE-three inhibition (Fig 4a). Similar but not significant changes in cell motility were observed in PANC-1 cells inhibited at CRE-three (G 7-8) (Fig. 4d). These results suggest a CRISPRi perturbation of CREs harbouring NCMs in the third intron of *MIR100HG* can decrease the migration ability in PANC-1 cells.

### Mutation occurrence of functional CREs in other solid cancers

Lastly, we explored the NCM burden of our top five prioritised regions (obtained from the first approach) in other cancers. We analysed the mutational frequency of these CRE-associated loci in seven other solid tumours using SSM data from the ICGC in oesophageal (ESAD), liver (LIHC), breast (BRCA-UK), ovarian (OV), prostate (PRAD-CA, PRAD-UK), colorectal (COAD) and gastric cancer (STAD) cohorts. The *HOTAIR*/*HOXC* CRE had the highest mutation frequency of NCMs across oesophageal (16.6%), liver (13.2%), prostate (7.5%) and ovarian (19.4%) cancers along with PDAC (5-12%, Extended Data Fig. 10). However, a low mutation frequency was observed in gastric, breast and colorectal cancers below 2%. The *FOXA1* CRE was predominately mutated in prostate cancer at an incidence of ∼16%, followed by liver, ovarian and oesophageal cancers at a frequency of ∼5%, higher than that observed in PDAC (2%). Interestingly, this regulatory region and NCMs have been recently reported in prostate cancer and are correlated with decreases in *FOXA1* expression and cell growth^81^. For the *MIR100HG* CRE, oesophageal and prostate cancer (UK cohort) showed the highest incidence at 14.2% and 5.7%, respectively, and liver and ovarian cancers showed a similar mutational incidence to the PDAC cohorts (2-3%). Other cancer types, such as breast, gastric and colorectal, had a very low to no mutational burden within this *MIR100HG* CRE (Extended Data Fig. 10c). The *FOXP1* CRE had the highest mutation frequencies in the liver, oesophageal and ovarian cancers (6-8%), but the *NFEL2* CRE generally had a much lower mutation frequency across all cancers, with a mutation burden of 2-3% in liver and oesophageal cancers, similar to that in PDAC. Our results suggest that several CREs identified in this study were also frequently mutated in other cancers. NCMs within these CREs may also play a functional role in contributing to these malignancies, as already documented in prostate cancer^81^.

## Discussion

Our study combines a computational discovery strategy and experimental follow-up to assess the functional significance of NCMs associated with PDAC-specific CREs. We leverage NCMs from PDAC SSM data derived from the ICGC ^7^ and integrate with PDAC-specific CREs marked by H3K27ac in seven PDAC cell lines and two patient-derived organoid samples. Previous investigations have often relied on consensus regulatory regions defined by ENCODE cell lines or the Ensembl Regulatory Build ^82^, this is likely to miss many enhancers which regulate genes in a highly cell and tissue specific manner ^83^. Our PDAC consensus peaks have incorporated high- and low-grade cell lines and patient derived organoids accounting for the tissue and stage specificity of regulatory elements associated with PDAC biology^21^.

The non-coding genome comprises a diverse spectrum of elements, and the mutational patterns and consequences are highly heterogeneous, rendering one approach ineffective across all regions of the non-coding genome ^84,85^. Thus, our pipeline incorporates an approach that directly estimates the functional consequence (i.e. deleteriousness) of each NCM and another that detects recurrently mutated CREs taking into consideration confounders such as replication timing and background mutation rates. Hence our combined approach identified a comprehensive, robust set of CREs subject to PDAC-relevant biological processes for *in vitro* validation.

High-throughput enhancer reporter assays are a powerful approach to screen the regulatory activity of a large number of NCMs in parallel ^40,43,86,87^. Our STARR-seq data highlighted 43 NCMs from PDAC patients showing significant gene reporter activity in the PANC-1 cell line. Interestingly, we observed the largest number of NCMs upstream of the *BNC1* promoter, resulting in a significant increase in STARR-seq reporter gene expression (Fig. 2d). Assessing the GEP of patients with these NCMs compared to those without NCMs demonstrated significant expression changes in more distal genes *BTBD1*^46^, *FAM103A1*^47^ and *TM6SF1*^48^. These DE genes were also associated with poorer overall outcomes in PDAC patients with higher expression (in *BTBD1* and *FAM103A1* genes) and lower expression for *TM6SF1* expressing patients (data not shown). Additional interesting candidates, such as NCMs proximal to cancer and PDAC-related TF *TBX3*^34^ and NCMs in the intron of *FOXP1*^88^, would be interesting and relevant candidates for future studies.

We identified significant CREs harbouring NCMs at the *MIR100HG* introns using both computational approaches, highlighting its importance for further functional validation. Previously, the transcription of *MIR100HG* has been linked to βT GexFp-ression/induction through SMAD2/3 binding sites in PDAC cell lines and *in vivo* studies leading to the release of its hosted miRs, including the oncogenic miR-100 and miR-125b-1 ^25,54^. The CRISPRi-based perturbation of *cis*-regions harbouring the most significant NCMs in the third intron of MIR100HG (identified using luciferase or high-throughput STARR-seq experiments) led to a down-regulation of *MIR100HG* expression and, in turn, cell mobility (Fig. 4c). This was correlated with a significant downregulation in critical PDAC related pathways included KRAS, P53, TGF-β and TNFα signalling^72,73^. Although not tested here, the direct targeting of these *cis*-regions leading to a down-regulation of *MIR100HG* transcription may inhibit the release of its hosted oncogenic miRs, as previously reported^25,54^.

Applying 4D genome interaction data ^62^, we observed looping of our targeted CRE-two with the promoter of proximal *EGFR-related* gene, *UBASH3B*^66^ and CRE-three with the promoter of the RhoA regulating GEF protein *ARHGEF12*^67^. Using RT-qPCR, we demonstrated CRE-two had the ability to downregulate *UBASH3B* expression, and CRE-three inhibition led to the significant reduction of *ARHGEF12*. These putative interactions may contribute to the downregulation of core pathways revealed by the GSEA analysis, as seen by the downregulation of EGFR signatures upon CRE-two inhibition ^66^ and MYC-target downregulation with CRE-three inhibition ^89^. This is the first report to our knowledge of NCMs in the introns of the lncRNA *MIR100HG* and the suggestion of *cis* genes other than MIR100HG being altered in expression ^25,54^. Considering a large number of transcripts MIR100HG has, further assessment of these CREs and NCMs on splicing would be important.

Genetic changes are critical for PDAC initiation, and up until recently, with the clinically available KRAS ^G12C^ inhibitor (AMG 510) ^90^ and the preclinical development of the KRAS^G12D^ inhibitor MRTX1133 ^91^, core mutated genes are largely undruggable. The reversibility of epigenetic changes allows the opportunity for therapeutic targeting. Previously in prostate cancer cells, the silencing of MIR100HG has led to the sensitisation to cytotoxic drugs^54^. We have shown here that perturbation of MIR100HG-associated CREs has collectively led to the downregulation of multiple core signalling pathways, including those previously not implicated in MIR100HG disruption, such as KRAS and TNFa signalling ^25,54^. In addition to the above considerations of this study, further investigation into the therapeutic potential of targeting this enhancer cluster rich in CREs and NCMs would be the next step.

We have limited this study to active enhancers widely reported to be marked by H3K27ac and H3K4me1^19^. However, we observed NCMs located outside of PDAC-associated CREs to have high functional predictive scores, suggesting they may lead to a gain/loss in functional activity at the gene level (Extended Data Fig. 2b). Moreover, use of H3K27ac alone to predict active enhancers may be too simplistic as many enhancers are marked with H4K16ac and H3K122ac but lack H3K27ac ^92,93^, suggesting many more CRE associated NCMs may be missed here. We have demonstrated the enhancer function for the *MIR100HG* locus harbouring PDAC-specific NCMs. However, further work is needed to demonstrate the pathogenic role of other NCMs identified in PDAC. Overall, our work identified and validated functional CREs and associated NCMs that may contribute to PDAC tumourigenesis and we have demonstrated a systematic framework to study *cis*-regulatory mutations in other human diseases.

## Methods

### Data acquisition

Data from the International Cancer Genome Consortium were downloaded from the ICGC portal (https://dcc.icgc.org/) release 27^7^. This data included simple somatic mutation (SSM) data for pancreatic ductal adenocarcinoma samples from the PACA-AU and PACA-CA cohorts. Clinical data, array-based expression (EXP-A from the PACA-AU cohort) and sequencing-based gene expression data (EXP-S from the PACA-CA cohort) were also downloaded. Gene Expression Omnibus (GEO) acquired datasets GSE64560 ^21^ and GSE99311^22^ were used to obtain ChIP-seq data to identify active enhancer-associated regions of the genome (H3K27ac and H3K4me1) based on seven PDAC cell lines and two patient-derived organoid samples. Additional marks were used to annotate further putative promoters (H3K4me3) and repressive domains (H3K9me3, H3K27me3).

### ICGC data processing

Downloaded SSMs were annotated and filtered using Annovar tools, retaining only those residing in non-coding elements (i.e., intergenic, intronic, synonymous and UTR) ^94^. Annovar ‘filter-based’ annotation method with packages: hg19_avsnp147, hg19_snp138, hg19_cytoBand, hg19_dbnsfp30a, hg19_ensGeneMrna was used. Available raw array-based expression (EXP-A) data was retrieved for 269 out of 391 patients from the AU cohort and normalised. Raw RNA-seq data for 234 out of 268 patients from the CA cohort were also downloaded. Quality-checked sequencing reads were aligned to build hg38 of the human genome using Hisat2 (version 2-2.1.0) ^95^ and annotated using Gencode release 27 hg38 ^96^. Read counts were estimated for each gene in all samples using HTSeq ^97^. Counts were normalised and transformed to log _2_-counts per million (log _2_CPM) using Voom (Limma package by BioConductor) ^98^. Log_2_CPM counts were then used as a measurement of gene expression.

### ChIP-seq data processing and manipulation

Raw sequencing reads in fastq files were extracted from GEO, and checked for quality using FastQC (version 0.11.5) ^99^. Where adaptors were present, sequences were trimmed using Trimmomatic tools ^100^. Subsequent reads were aligned to the human reference genome (hg38) using Bowtie2 (verison 2/2.3.0) with default parameters ^101^, and duplicate reads were marked with Picard (MarkDuplicates) ^102^ and removed using SAMtools ‘rmdup’ ^103^. Uniquely aligned reads were downsampled between ChIP-seq samples and input control pairs to avoid read yield bias. Genome-wide narrow peaks were called for H3K27ac and transcription (TF) samples, and broad peaks for H3K4me1, H3K4me3 and H3K9me3 samples against the input control using MACS2 (version 2.1.0) default settings where data was available^104^. Peaks were further filtered for quality, preserving peaks with a Q-value of E-10. Subsequent BedGraph file outputs from MACS2 were converted to BigWig files using the UCSC binary tool, BedGraphToBigWig. H3K27ac peaks located with an inter-peak distance of 2,000bp to other PDAC cell line H3K27ac regions, were merged using the ‘merge’ function from Bedtools (version 2.26.0) to produce one consensus H3K27ac region across all samples. H3K27ac peak co-ordinates were ‘lifted’ over to hg19 using the UCSC command line tool ‘liftOver’ to overlap with SSMs. H3K27ac regions harbouring non-coding mutations affecting >2% of the patient cohort were retained for further analysis (≥8 NCMs in ≥8 patients).

### The identification of putative functional mutations using approach one (non-coding annotation/IW-scoring and LINSIGHT algorithms)

SSMs from filtered and merged H3K27ac peaks were subjected to functional testing and filtering using the IW-scoring algorithm ^23^. The workflow for the identification of novel variants was utilised, excluding the use of GWAVA scores (for known variants). The median IW-functional score for all mutations within each H3K27ac consensus region was calculated. H3K27ac regions with a median IW-score of two or above were retained for further analysis. In addition, IW-scores of NCMs residing outside (H3K27ac negative) the top candidate H3K27ac consensus regions (∼1kb) were obtained and compared to those of H3K27ac associated NCMs. The top candidate regions were also validated using the LINSIGHT algorithm. LINSIGHT scores were extracted as previously described ^31^. The scores based on the likelihood of deleterious fitness consequences were extracted and used to compare NCMs located inside our consensus peak regions and NCMs located nearby outside peak regions (H3K27ac negative). An unpaired Wilcoxon signed rank test was used for all statistical significance testing.

### The identification of putative functional mutations using approach two (LARVA algorithm)

To identify recurrently mutated regions (within H3K27ac consensus peaks) more than expected to nearby background regions, we used the algorithm LARVA ^32^. This algorithm considers sample-specific mutation rates, heterogeneity and replication timing, as previously described ^32^. NCMs that fell into blacklist regions were first removed, and the remaining NCMs overlapped with our H3K27ac consensus regions. Three models were used to calculate the mutation rate expected based on the stochastic background mutations. The first and second model calculates the number of local mutations within a given annotated region and estimates the probability of observing a mutation in each position. The *p*-value was drawn from a β-distribution, taking the average mutation rate and the over-dispersion, respectively into consideration. The third model considers the average replication timing within each H3K27ac element, a confounding genomic feature that would affect the background mutation rate^85^. For this, replication timing data from seven different cell lines were retrieved from ENCODE and the average timing per region calculated across all cell lines (HepG2, MCF-7, GM12878, K562, BJ, IMR-90 and SK-N-SH GSE34399)^105^. *P*-values were adjusted with the Benjamini-Hochberg method across all three models. We prioritised those significant H3K27ac regions with a *q* value of <0.01.

### Luciferase reporter assays

Sequences surrounding NCMs of interest (∼2kb total) were amplified using specific primers (Extended Data Table S4). Mutations were introduced with site-directed mutagenesis (QuikChange II Site-Directed Mutagenesis Agilent) as per the manufacturer’s instructions and checked using Sanger Sequencing and correct regions cloned into the pGL2 vector upstream of the SV40 promoter. Thirty-five thousand cells (HEK293T and PANC-1) were plated 24-hours before transfection in a 24-well plate with either 100ng WT or MUT pGL2 plasmids (Promega Cat E1631) and 5ng of Renilla luciferase control (Promega Cat E2231). Luciferase activity was measured 48-hours post-transfection with the Dual-Luciferase Reporter Assay System (Promega Cat E1910). Overall activity was calculated by taking a ratio of the Firefly over the Renilla expression control vector. The background signal was quantified using un-transfected cells and subtracted from readings. An unpaired *t-test* was used to obtain statistical significance between wild-type (WT) and mutant (MUT) luciferase activity.

### STARR-seq library design and cloning of candidate *cis*-regions into the STARR-seq plasmid

The STARR-seq library consisted of 6,080 constructs representing 587 candidate mutations, corresponding WT sequences and 210 controls. Constructs were represented in a 194bp sequence context, flanked by a 15bp linker region for adaptor ligation and amplification (Extended Data S4). One hundred and ten positive controls were selected from super-enhancers previously reported in PDAC ^106^ and additional regions from the super-enhancer database (SEdb) ^107^. Putative negative controls were selected from gene deserts lacking H3K27ac and H3K4me1 marks in PDAC cell lines. A unique 6bp barcode was placed between the 5’ 15bp linker and the candidate sequence to allow the differentiation between WT, MUT and control (CTRL) sequences, resulting in a final construct of 230bp. To understand the activity of mutations in different genomic contexts and maximise the chance of capturing regulatory activity, each mutation was represented in the library five times, shifting the genomic context of the sequences 10bp and 20bp left and right from the middle of the construct, thereby representing the mutation in left_20bp, left_10bp, centre, right_10bp and right_20bp positions. The synthetic oligonucleotide library was amplified and cloned as previously described^44,108^. Briefly, 5ng of the STARR-seq library was amplified, and vector homology arms were added to either side of the construct. The second generation hSTARR-seq ORI plasmid (Addgene: #99296) was digested with Sall-HF and Agel-HF restriction enzymes, and the amplified library was cloned into the 3-UTR of the vector. Ligations (X5 reactions) were transformed by electroporation into MegaX DH10B™ T1R Electrocomp™ Cells (Invitrogen), and reactions pooled. The plasmid pool was extracted using the ZymoPURE Giga prep kit according to the manufacturer’s instructions. To check the quality and overall representation of the library, sequence inserts were amplified from the STARR-seq plasmid using Illumina-compatible index primers (Extended Data S4). STARR-seq libraries were sequenced using 2 x 150bp chemistry on an Illumina Novaseq 6000 by Novogene ltd.

### STARR-seq oligo-pool quality check

Paired end reads were merged into single amplicons using the USEARCH fastq_mergepairs command^109^. Merged reads were aligned back to the expected oligo library using BWA MEM with default parameters, penalising soft-clipping of alignment ends (-L80) ^110^. GATK DepthofCoverage (version 3) was used to determine the sequencing depth per nucleotide and construct ^111^. Of the 6,082 constructs sequenced, 98.63% had a minimum coverage of 30X, with both WT and MUT sequences represented. To identify sequencing errors, the Samtools ‘mpileup’ function was run on aligned reads and the oligo reference library to obtain read counts for each nucleotide position^103^. Subsequent mpile up files were run with the VarScan2 package and ‘mpileup2cns’ parameters to identify sequencing errors^103,112^.

### Transfection, RNA isolation and cDNA synthesis

Two million PANC-1 cells were plated per 10cm dish (5 dishes per biological replicate) for 24 hours. Plasmid libraries (14 μg per plate) were transfected using lipofectamine 3000 as per manufacturer instructions. To monitor transfection efficiency, one 10cm dish was co-transfected with 2.8μg of pmaxGFP plasmid (Lonza). Immediately post-transfection, the interferon inhibitors C16 and BX-795 were added to each plate at a final concentration of 1μM (per inhibitor), as previously described^113,114^. Cells were incubated at 37°c for 16 hours before harvesting and counting. 1/10 ^th^ of the cells were retained for plasmid DNA, and the remaining cells were for RNA extraction. For RNA, cells were homogenised with the Qiagen Qiashredder and total RNA was extracted using the Qiagen mini-RNA extraction kit as per the manufacturer’s instructions. Poly-(A)^+^ RNA were isolated using Dynabead^T^s^M^ oligo(dT)25 followed by DNase treatment with TurboDNase (Invitrogen). Samples were purified with RNA cleanupXP beads as previously described ^108^. cDNA synthesis was carried out using SuperScript III and a gene-specific primer (Extended Data S4). cDNA was purified with 1.4X AMpureXP beads (and for subsequent steps described below). A second-strand synthesis reaction was followed by purification. Using a P7-specific primer (Extended Data Table S4) UMI’s were added to cDNA (in 5 reactions) with Kapa 2x HiFi HotStart ReadyMix (Kapa Biosystems). Reactions were pooled and purified. Junction PCR was used to amplify reporter-specific transcripts for 16 cycles and thereafter purified. For the final library preparation, Illumina sequencing primers were used in cDNA samples for 8-14 cycles followed by purification with 1.2X of AMPure SPRI beads (Extended Data S4).

To obtain the DNA input library, STARR-seq plasmids were isolated from PANC-1 cells using the Monarch plasmid miniprep kit, as per the manufacturer’s instructions. One hundred nanograms of DNA were amplified using Illumina-compatible index primers as described above. The DNA plasmid and RNA-derived libraries were sequenced using the 150-cycle paired-end V3 chemistry reagents and run on a Miseq.

### Processing and analysis of STARR-seq screen

Paired-end reads were processed with CutAdapt to remove residual sequencing adaptors and STARR-seq vector linkers^115^. Reads were split based on the 6 bp barcodes WT, MUT and CTRL into separate files. Barcodes were removed, and sequences aligned to the human reference genome (hg19) using BWA MEM with default parameters ^110^. Aligned BAM files were converted to BAMPE format using the bedtools function ‘bamtobed’, and properly paired reads were extracted for further analysis ^116^. The Bedtools ‘intersect’ function was used to overlap reads with the expected design oligo library and obtain raw read counts. Samples were deduplicated based on UMI’s with a custom-made Perl script. A minimum of three unique UMI’s were required for a construct to be counted. Deduplicated counts were normalised to the total number of reads in the sample and then multiplied by 1M to obtain the number of transcripts per million. The relative abundance of each construct transcribed was calculated by dividing the observed RNA output by the DNA input, indicating the relative activity of each WT, MUT and CTRL construct. To compare the transcriptional activity of single oligos between WT vs MUT and negative vs positive CTRLs, an unpaired t-test was used. To compare the transcriptional activity at the mutation level across the five sliding windows (WT vs MUT), a Mann-Whitney U statistical test was used.

### CRISPRi guide RNA design and cloning

For CRISPRi, guide RNAs were selected from the UCSC genome browser ‘CRISPR Tracks’, selecting guides as close to mutations as possible with a minimum of two guides per *cis*-region (Extended Data Table S4). Potential off-target effects were assessed using the MIT specificity score, selecting guides with a score above 70%^117^. Homology arm sequences were added to each guide to clone into the pU6-sgRNA EF1Alpha-puro-T2A-BFP expression plasmid at the BstXI-BlpI31 digested site. gRNA oligos were phosphorylated, annealed and cloned into pU6-sgRNA EF1Alpha-puro-T2A-BFP expression plasmid (Addgene #60955) as previously described ^118^. Inserts were verified with Sanger sequencing.

### Lentivirus transduction

Lentivirus was generated as previously described ^118^. Briefly, 4M cells were plated in a 10cm dish for 24-hours before transfecting HEK293T cells with 9ug of dCas9-mCherry-KRAB (Addgene #60954), 4ug of packing plasmids psPAX.2 and 2ug of the envelope vector pMD2.G diluted in OptiMEM medium and Trans-Ltl transfection reagent (Mirus). For the generation of gRNA lentivirus, 9ug of each cloned guide were transfected, and the virus was collected as described above. Twenty-four hours post-transfection, media was refreshed, and viral supernatant was collected at 48- and 72-hours post-transfection. Viral supernatants were centrifuged and filtered (45um). PANC-1 cells were transduced in a one-to-one dilution of the virus and growth medium supplemented with polybrene (5ug/ml). Three days post-transduction, mCherry positive cells were sorted by FACs, selecting the top 50% of positive cells based on the overall mCherry signal. PANC-1 dCa9/KRAB expressing cells were plated in 24-well dishes for 24-hours before transducing cells with lentiviral supernatant from multiple guides (as indicated in Fig. 3b). At 24-hours post-infection, the medium was replaced, and cells were selected with 2ug/ml of puromycin for 72-hours. Cells were harvested, and the effect on the expression of MIR100HG, UBASH3B and ARHGEF12 was assessed using qPCR and subsequent RNA-sequencing (Extended Data Table S4).

### qPCR

RNA was extracted and DNase I treated using the Qiagen mini-RNA extraction kit according to manufacturer instructions. cDNA was synthesised from 1ug of DNase treated RNA using the LunaScript® RT SuperMix (NEB), according to the manufacturer’s protocols. We performed qPCR on a StepOneTM Real-Time PCR System with the Luna® Universal qPCR Master Mix (NEB). Gene specific primers are outlined in Extended Data Table S4.

### RNA-seq data generation and analysis

500 ng of total RNA was used to enrich mRNA using an oligo dT-based mRNA isolation module (NEB Cat number E7490L). RNA sequencing libraries were prepared by NEBNext Ultra II Directional RNA Library Prep Kit for Illumina (NEB catalogue number E7760S). Libraries were sequenced as 150 bp paired-end reads using a Novaseq 6000. After the quality check and trimming, reads were aligned to the reference genome hg38 using STAR v2.7.9a ^119^, followed by the gene count quantification using RSEM ^120^ based on the Ensembl gene annotation GRCh38.p13 Release 105. Genes with low mapped read across all samples were removed. The normalised RPKM (Reads per kilobase of transcript per Million reads mapped) expression values for all filtered genes across samples were subsequently derived and used for the differential expression (DE) analysis. The DE analysis was performed using Limma^121^, comparing each CRISPRi perturbation group (G 5-6 and G 7-8) to the dCa9/KRAB control group respectively. The significant DE genes were identified using a threshold of FDR<0.05 and absolute log_2_FC>1. GSEA ^68^ was then performed based on the Limma output against gene sets curated in MSigDB hallmark^69^ and oncogenic signature gene sets, to identify dysregulated gene activities in the CRISPRi group relative to the control.

### Cell migration assays

Approximately four thousand dCas9/KRAB expressing PANC-1 cells transduced with lentiviral gRNA combinations were seeded into 96-well plates. Cells were scratch wounded using a 20ul pipette tip. Cells were washed with PBS to remove cell debris, and phase-contrast images were taken at 0-, 24- and 48-hours at three specific wound sites per well using a Leica microscope with an X4 objective. The ability of the cells to migrate and close the wound area was evaluated by comparing the pixels of the open wound region at each time point using image J (MRI wound healing plugin) ^122^. An unpaired t-test was used to compare each treated time point to the negative control.

## Data availability

The RNA-seq data for the CRISPRi perturbation of MIR100HG enhancer regions has been deposited to the Gene Expression Omnibus under the accession number of GSE229499. ChIP-seq data were available under GSE64560 and GSE99311. Mutation and expression data of PDAC patients were downloaded from the ICGC data portal. The STARR-seq data and all scripts to analyse the data can be requested and obtained by contacting the corresponding authors.

## Acknowledgement

The work was funded by the Medical Research Council Doctoral Training Partnership (DTP) PhD programme to QMUL and University of Southampton, and by Academy of Medical Sciences Springboard Award (SBF003\1025 to J.W.). The authors acknowledge support from the Cancer Research UK City of London Major Centre core funding support to Barts Cancer Institute (C16420/A18066), and the Accelerator Award Program funded by Cancer Research UK (C355/A26819) and FC AECC and AIRC. Medical Research Council UKRI/MRC grant (MR/T000783/1) (MMP, DP) Barts charity small grant (MGU0475) (MMP).

## Author contributions

J.W. conceived the study. J.W., M.M.P., and M.B.P. designed the study and developed the methodology. M.B.P., and J.W. performed the computational discovery analysis. M.B.P., A. R.-M., J.H., J.F., and M.M.P. performed the Luciferase reporter assay experiments. M.B.P., D.P., and M.M.P. performed the STARR-seq experiments. M.B.P., S.S.A., and J.W. performed the STARR-seq data analysis. M.B.P., E.M., and J.W. performed the RNA-seq analysis. M.B.P., H.K., and M.M.P. performed the in vitro functional assays. J.W., and M.M.P. supervised the study and acquired the funding. M.B.P., J.W., and M.M.P. interpreted the data and wrote the manuscript. All authors reviewed the manuscript and approved the final version of the manuscript.

---

**Extended Data Fig. 1. Prioritised CREs using the first** *in silico* **approach.** (**a**) Australia (AU) and (**b**) Canada (CA) cohorts NCMs were submitted to the IW-scoring algorithm. Each dot denotes a CRE, it’s combined median IW-score, across all chromosomes. The horizontal dotted line indicates the median IW-score threshold (p=0.1). The prioritised CREs with an IW-median score ≥2 are labelled by the nearest proximal gene for each cohort.

**Extended Data Fig. 2. An independent validation of the top prioritised CREs using LINSIGHT. a**. Boxplots of LINSIGHT scores in H3K27ac-associated NCMs (inside peaks) in comparison to NCMs in nearby H3K27ac negative regions (outside peak) for the top 5 CREs identified by the first approach. **b**. LINSIGHT scores and location of NCMs inside (green) vs outside (black) H3K27ac peaks.

**Extended Data Fig. 3. Identification of two putative CREs within the** *MIR100HG* **enhancer cluster.** The MIR100HG enhancer cluster was the only shared element between the two in-silico approaches. One CRE was identified by the first approach based on IW-scoring and LINSIGHT validation, and the other was identified by the second approach based on the LARVA algorithm. Genome browser tracks (hg19) of the H3K27ac peaks across the 7 PDAC cell lines and 2 patient-derived organoids are shown. The MIR100HG-hosted microRNAs and associated gene *BLID* are indicated.

**Extended Data Fig. 4. First** *in silico* **approach prioritises a significant CRE in the third intron of the TF** *FOXP1*. Genome browser tracks (hg19) presenting the H3K27ac peaks across the 7 PDAC cell lines and 2 patient-derived organoids and the annotation of the merged consensus peaks. The location of NCMs inside and outside H3K27ac consensus is indicated for each cohort.

**Extended Data Fig. 5. Quality analysis of the cloned STARR-seq plasmid library. a.** Length distribution of cloned oligo constructs. The percentage of each length is shown. The oligo construct library had 49% of oligos with the expected correct length, followed by 1-(28%) and 2-bp (10%) deletions. **b.** Depiction of the number of construct synthesis errors across the sequenced oligos. The error occurrence is shown along the base pair positions.

**Extended Data Fig. 6. Comparison of regulatory activities derived between Luciferase reporter assay and STARR-seq.** Boxplots comparing the Luciferase reporter assay activity and STARR-seq, demonstrating concurrent directional changes in enhancer activity at NCMs profiled by both techniques.

**Extended Data Fig. 7. BNC1 associated enhancer cluster and significant NCMs from the STARR-seq screen. a.** Genome browser (hg19) of the H3K27ac peaks across the 7 PDAC cell lines and 2 patient-derived organoids. The STARR-seq significant NCMs are indicated, with the vast majority residing upstream of the *BNC1* gene promoter. **b.** Boxplots showing the expression of genes within 1Mb of the *BNC1* CRE that have significant alterations (*BTBD1*, *FAM103A1* and *TM6SF1*) and the nearest proximal gene (*BNC1*), comparing mutant (MUT) and wildtype (WT) patient gene expression profiles (GEPs). The p-values were derived from the Wilcoxon rank sum test. **c**. Motif gain example for a gain-of-function NCM proximal to the *TBX3* gene (chr12:115067012:C>A). A binding motif for the TF JUN is created by this mutation. All TF binding predictions were carried out using MotifBreakR. **d.** Motif break example for one loss-of-function NCM located in the intron of *FOXP1* (chr3:71123616:G>T). The binding motif for ATF3 was disrupted by this mutation.

**Extended Data Fig. 8. The interaction of MIR100HG CREs with distal genes revealed by the 4D genome. a**. Genome browser (hg19) showing the H3K27ac signal for PANC-1 cells and the putative loop between CRE-2 and the promoter of *UBASH3B*. Putative loops were predicted using the interactions from the integrated method for predicting enhancer targets (IM-PET) and 4D Genome in PANC-1 cells. **b**. Genome browser (hg19) showing the H3K27ac signal for PANC-1 cells and the putative loop between CRE-3 and the promoter of *ARHGEF12*.

**Extended Data Fig. 9.** Boxplots comparing the gene expression profiles of TGF-β related genes *FGF1*, *KDM6B*, *LIF*, *PIK3CD*, *PXDC1* and *TAGLN* between G 5-6 / G 7-8 CRISPRi perturbations and the dCas9/KRAB control. The gene expression levels were measured by RNA-seq data, in the unit of log2 RPKM values.

**Extended Data Fig. 10. The mutational burden in the top five significant CREs identified in the first approach in other common solid tumours.** CREs overlaying genes *FOXA1*, *FOXP1*, *HOTAIR/HOXC* genes, *MIR100HG* and *NFEL2* were assessed for their mutational burden. **a.** Barplot showing the number of samples across the selected cancer cohorts. **b.** Barplot of the total number of mutations within each cohort. **c.** Barplots showing the frequency of NCMs identified within each CRE across each cancer cohort.

**Extended Data S1.** List of NCMs in significant CRE’s prioritised using the first *in-silico* approach in the AU and CA cohorts respectively.

**Extended Data S2.** Table showing the comparison of NCMs inside the top five significant CREs to NCMs located outside flanking H3K27ac negative regions. An unpaired Wilcoxon signed rank test was used to obtain p values.

**Extended Data S3.** List of all significant CREs prioritised by the LARVA algorithm and those significant CREs found to be in common between the AU and CA cohorts.

**Extended Data S4.** List of primers and CRISPRi guides.

**Extended Data S5.** List of top significant oligos as shown in Fig. 2e.

