## Supplementary Figures for "Non-coding mutations at enhancer clusters contribute to pancreatic ductal adenocarcinoma"

### Extended Data Figure 1

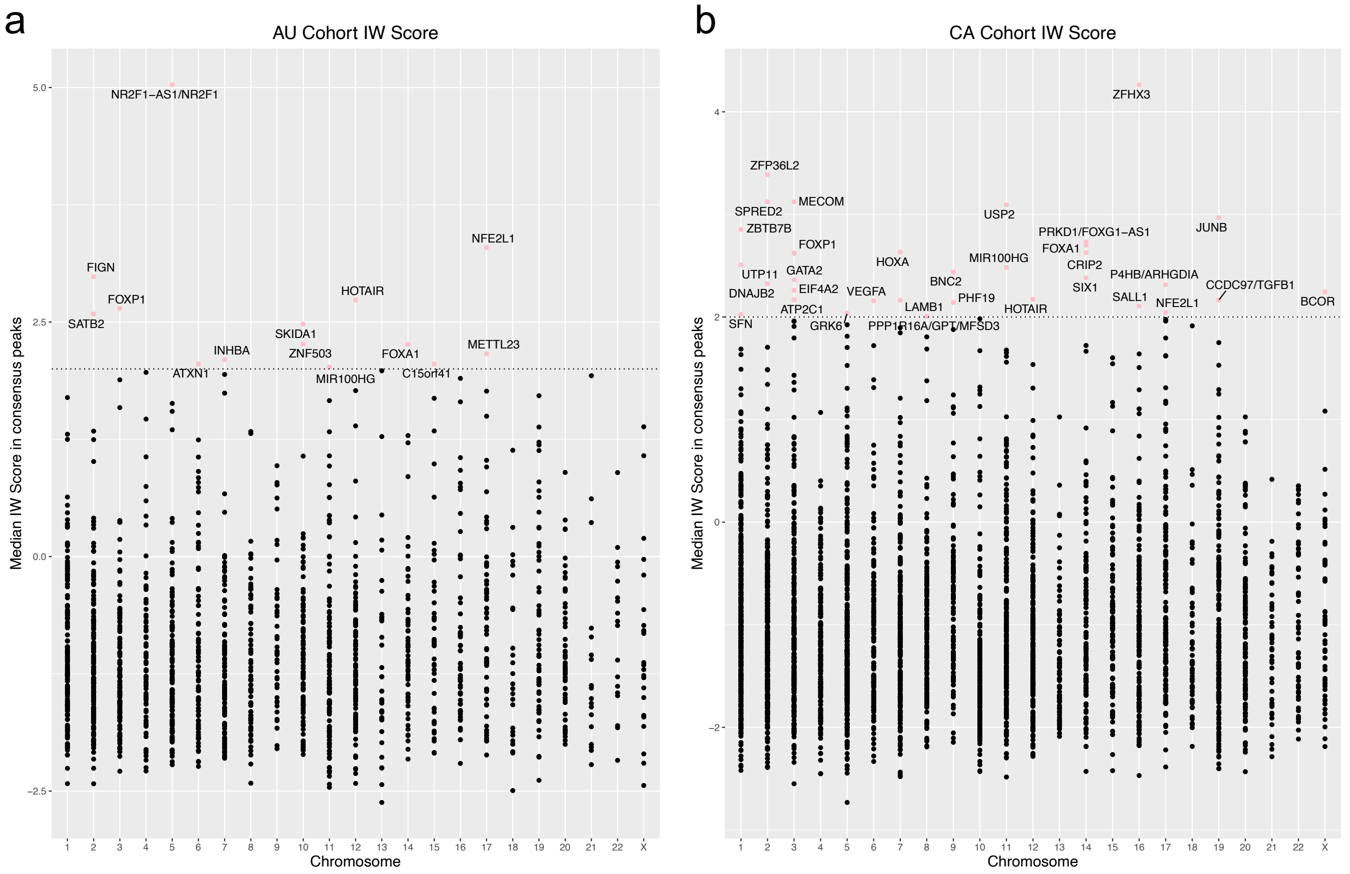

### Extended Data Figure 2

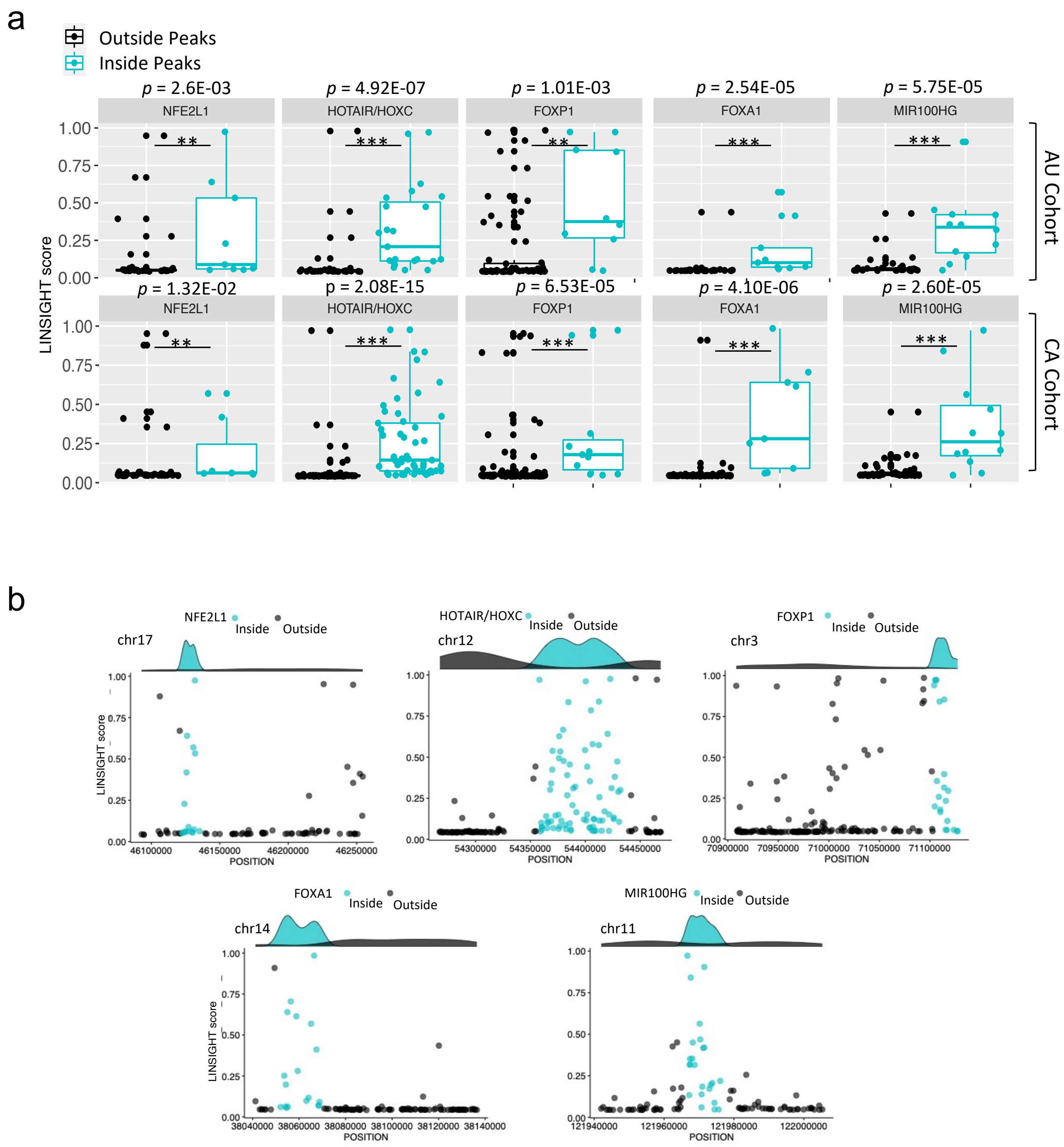

### Extended Data Figure 3

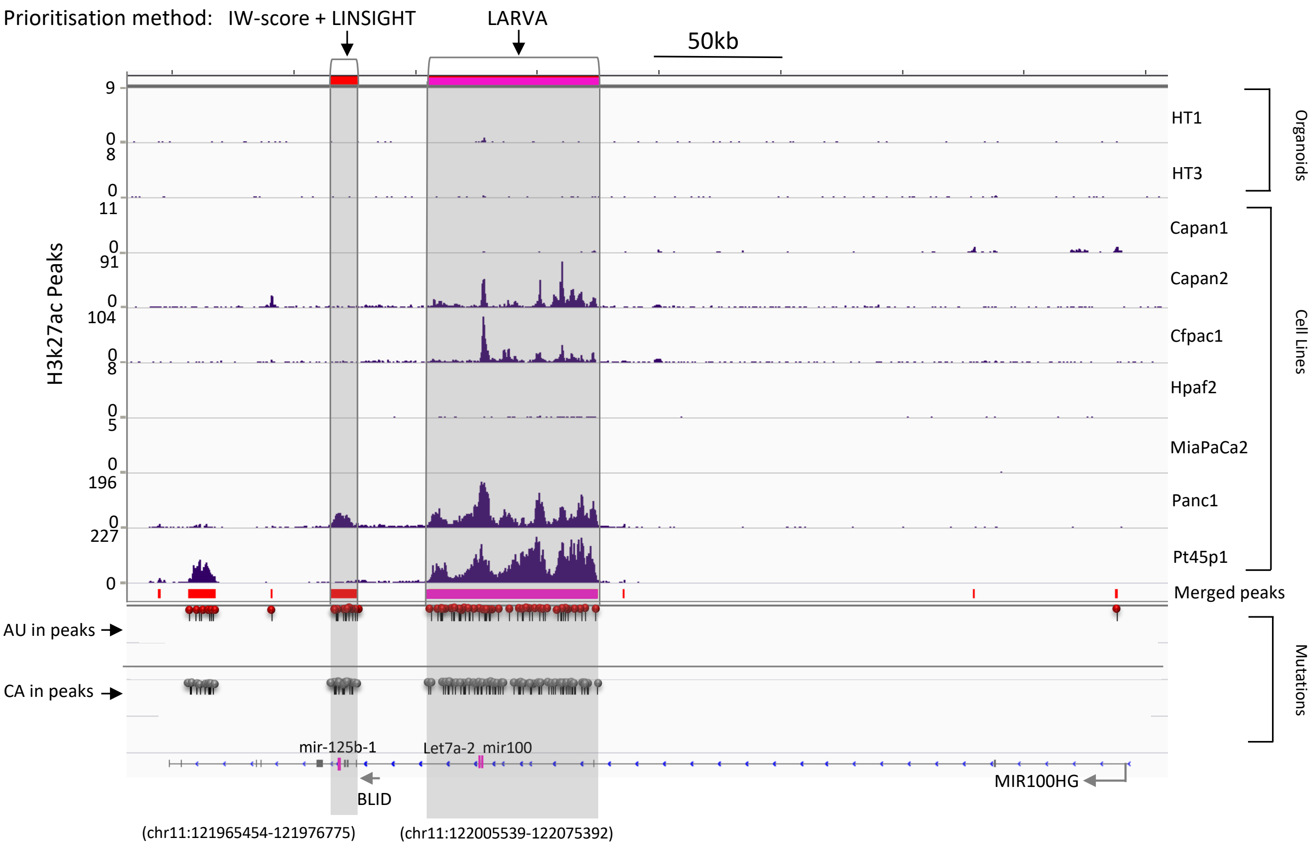

### Extended Data Figure 4

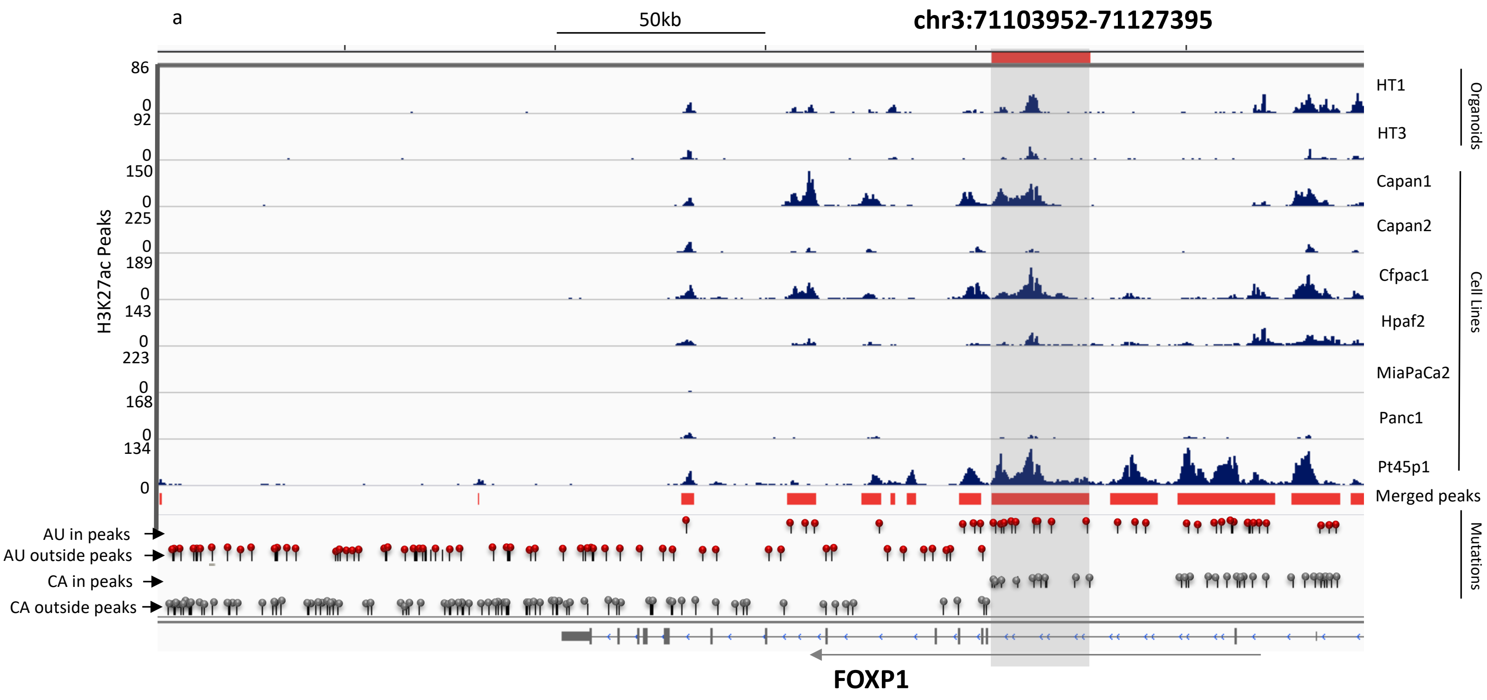

### Extended Data Figure 5

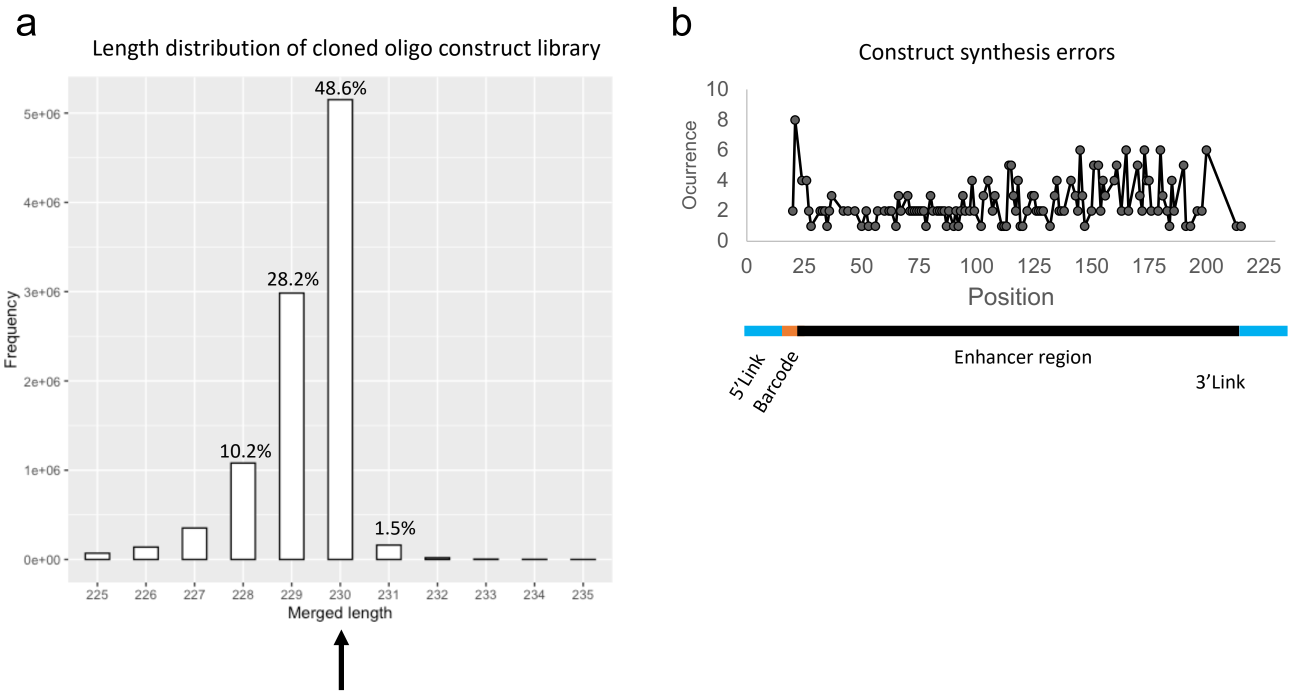

### Extended Data Figure 6

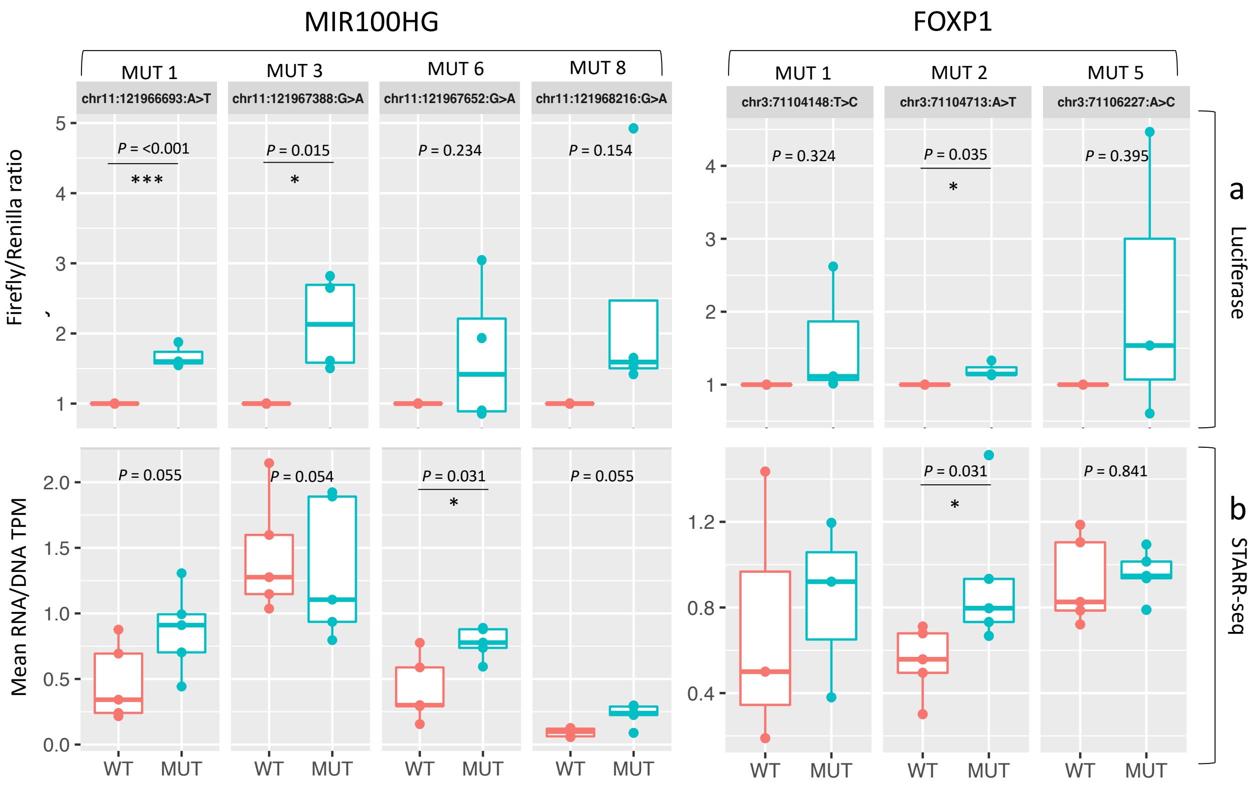

### Extended Data Figure 7

a

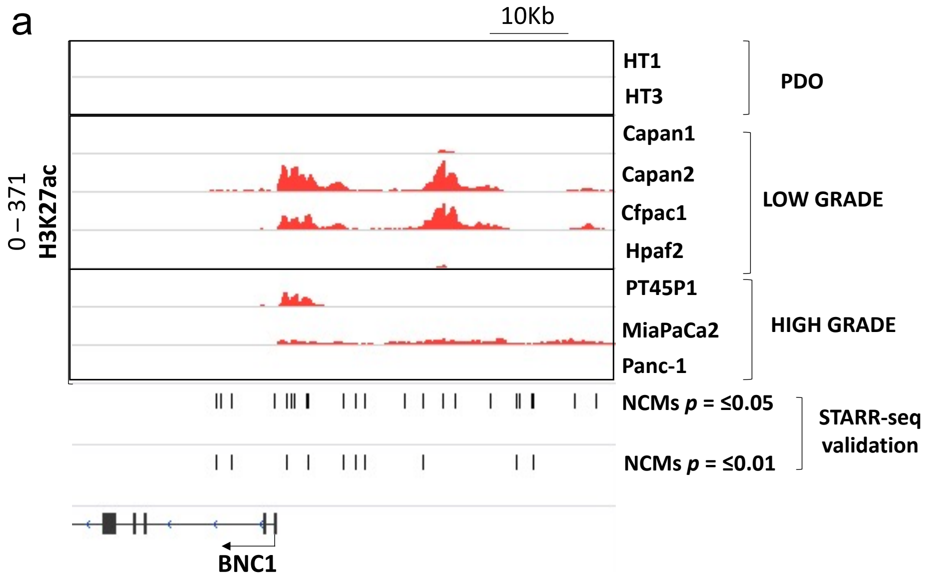


b

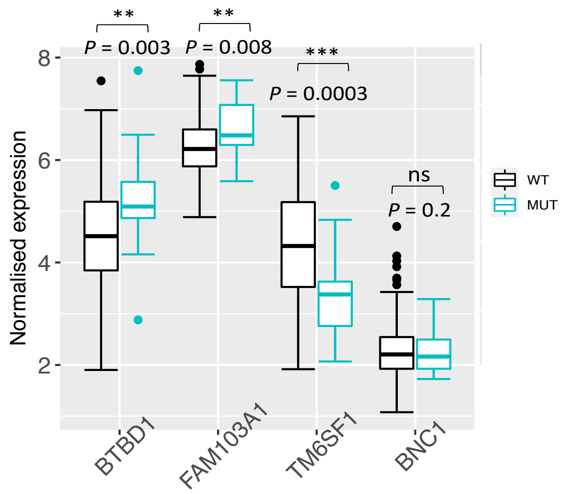


c

Motif Gain

JUN

chr12:115067012:C>A

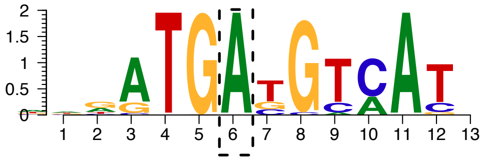

d

Motif Break

ATF3

chr3:71123616:G>T

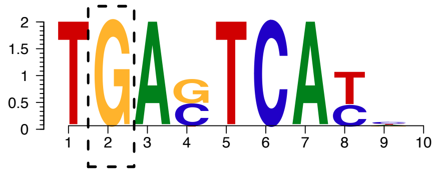

### Extended Data Figure 8

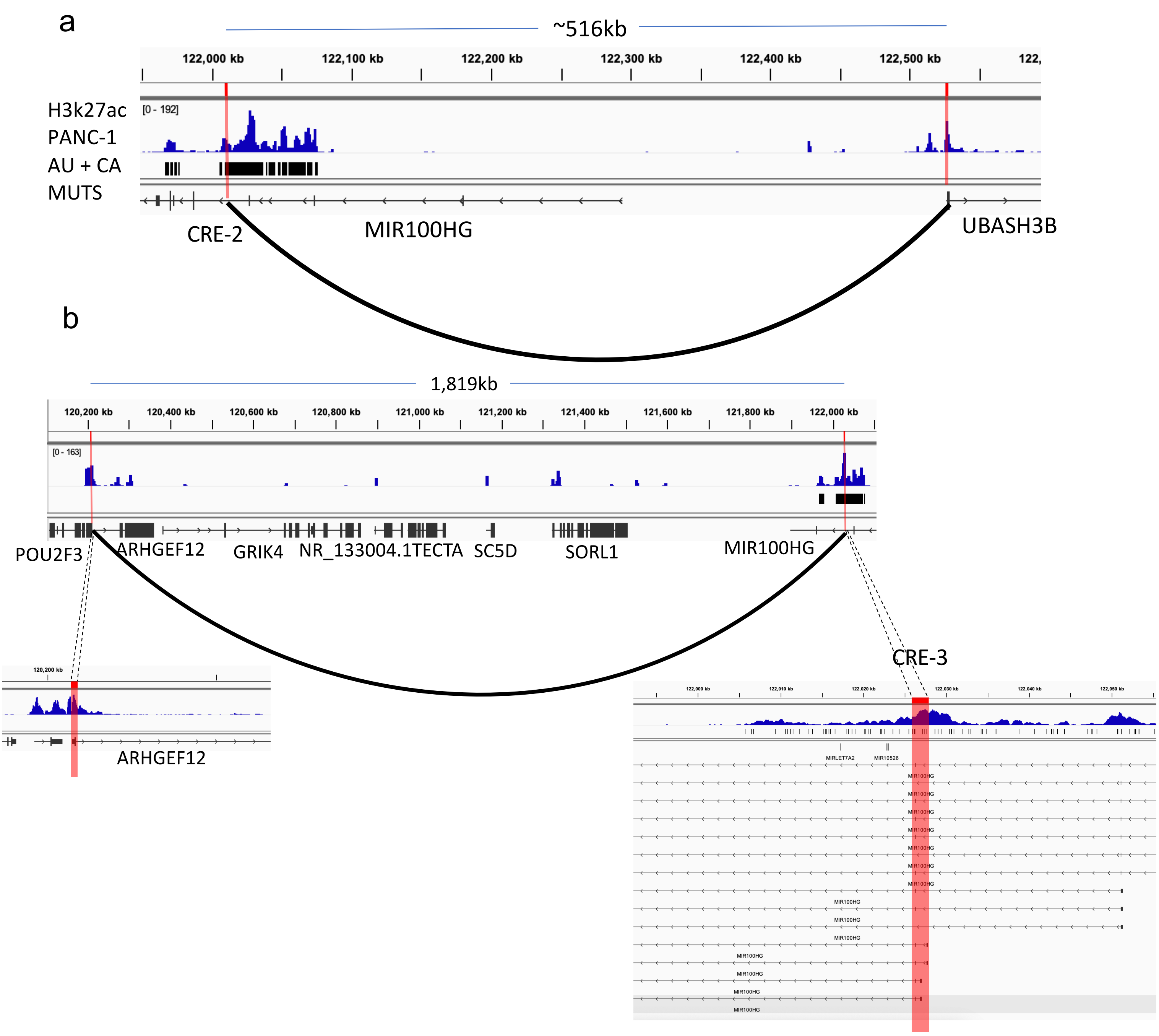

### Extended Data Figure 9

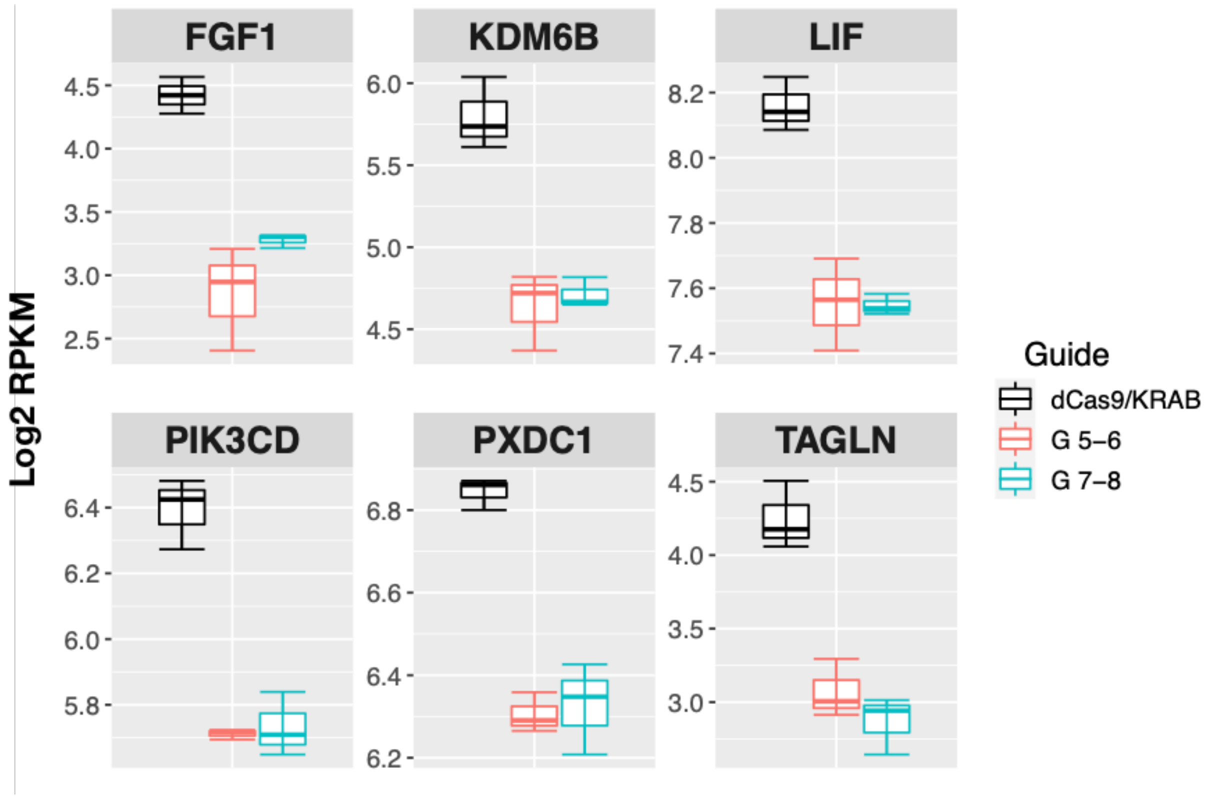

### Extended Data Figure 10

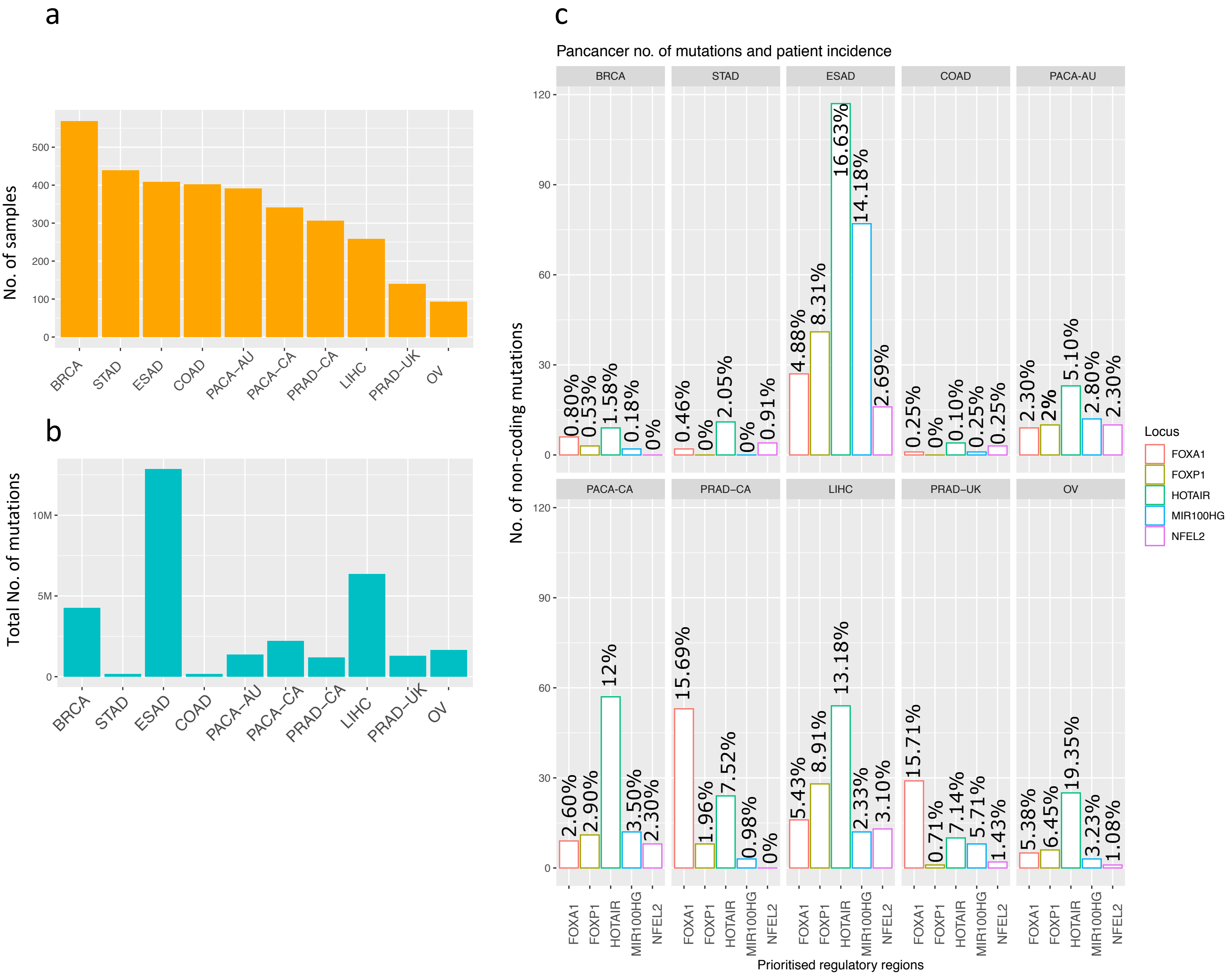
